# To self or to clone? Southern European woodland strawberry genotypes self-fertilize, whereas eastern European genotypes clone in a pollinator-free common garden

**DOI:** 10.64898/2026.03.30.715235

**Authors:** Carolina Diller, Ivan M. De-la-Cruz, Paul A. Egan, Timo Hytönen, Johan A. Stenberg

## Abstract

**Premise of study:** Under increasingly frequent pollinator-limited environments, plants need to rely on modes of reproductive assurance such as selfing and cloning. However, few studies investigate the interplay between selfing and cloning in plants that can do both. Here, we explore mechanisms determining the relative expression of selfing and cloning throughout the European distribution of the wild woodland strawberry (*Fragaria vesca*) under a pollinator-free environment.

**Methods:** We established an outdoor common garden with 121 woodland strawberry genotypes from across Europe and excluded them from pollinators. For each genotype, we recorded reproductive traits and performed hand-pollination treatments.

**Key Results:** We found a weak trade-off between cloning and selfing, driven by increased seed and fruit provisioning rather than flower production. The capacity to autonomously self-fertilize was determined by the lateral proximity of the anthers to the pistils (lateral herkogamy), but not by early inbreeding depression. Genotypes sampled at lower latitudes and altitudes were better at self-fertilizing and had smaller petals. The propensity to clone increased towards the east, where genotypes also had smaller petals, particularly at higher latitudes.

**Conclusion:** At the species level, we detected a trade-off between the propensity for clonal reproduction and the capacity for self-fertilization. At a continental scale, the capacity to self-fertilize varied along a north–south gradient, whereas clonal propensity varied along an east–west gradient. Our results suggest that these two modes of reproductive assurance may compensate for reduced pollinator attractiveness (smaller petals) in regions where each mode is most strongly expressed.

## INTRODUCTION

In the face of the current pollinator crisis, plants will increasingly need to rely on some mode of reproductive assurance (Potts et al., 2010; Marshman et al., 2019; Rodger et al., 2021). Pollen limitation, however, is not a new circumstance for plants and a wide variety of plant mating strategies exist to cope with it (Lloyd, 1992; Lloyd and Schoen, 1992; Goodwillie and Weber, 2018). Self-compatible plants, for instance, can autonomously self-fertilize if the flower morphology facilitates it (Kalisz et al., 2004). Clonal propagation, may also serve as a mode of reproductive assurance (Baker, 1955; Holsinger, 2000; Eckert et al., 2006; Vallejo-Marín et al., 2010) particularly in self-incompatible plants (Vallejo-Marín and O’Brien, 2007), but not exclusively. In fact, several plants can both self and clone. Studies on the mating system of plants that can both self and clone, predominantly revolve around the increased risk of geitonogamy (within-individual/genotype selfing) under clonal growth (Kron et al., 1993; Eckert, 2000; Albert et al., 2008; Marriage and Kelly, 2009; but see Van Drunen et al., 2015) (Fig. 1A). Yet without pollinators present, both outcrossing and geitonogamous selfing are low, and thus autonomous selfing and cloning serve as two possible modes of reproductive assurance (Fig. 1B). Are plants that are capable of both reproductive assurance strategies, employing them equivalently?

**Fig. 1.**
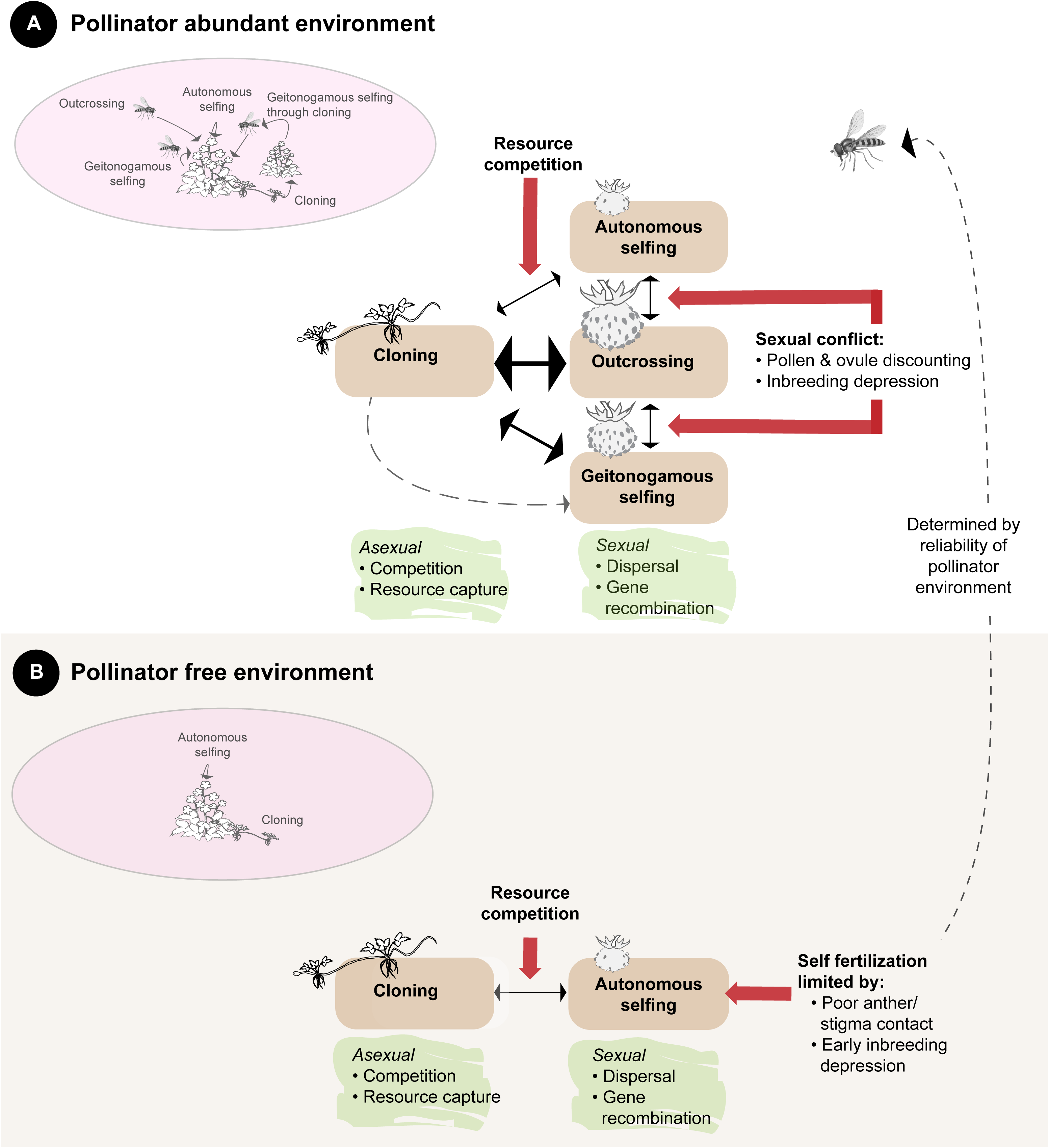
Mating system dynamics in a (A) pollinator abundant versus a (B) pollinator-free environment for a self-compatible plant that can both autonomously self-fertilize and clone. While the benefits of sexual and asexual reproduction do not change in a pollinator abundant vs. a pollinator-free environment (green background), the negative interactions do (red arrows). The width of the black arrows representing resource competition between asexual and sexual reproduction indicates the expected strength of this relationship (wider arrows denote stronger competition).

Dole (1992) recorded the differential expression of selfing and cloning across three species under experimentally induced pollen limitation. He found a positive relationship between selfing and cloning in the species with mixed mating system (i.e. species that can self and outcross). However, in the two species that were predominantly selfers, autofertility was higher than cloning. In their system, life history and habitat type seemed to be responsible for the differential allocation to selfing and cloning (Dole, 1992; Van Kleunen, 2007). Here, we consider an additional mechanism: how variation in the quality of selfed fruits and seeds affects the relationship with cloning.

Sexual and asexual reproduction may compete for resources, resulting in a trade-off. The strength of this trade-off may depend both on overall resource availability and on the magnitude of sexual reproductive investment, such as the number, size, and quality of fruits or seeds produced. Sexual output, in turn, is strongly influenced by the mating and breeding characteristics of the species and populations involved. For example, in mixed mating systems, selfing often produces smaller fruits or reduced seed set (Klatt et al., 2013) potentially resulting in a weaker trade-off with clonal reproduction (Fig. 1A). Lower reproductive output via selfing within mixed mating systems may arise from poor contact between anthers and stigmas (e.g. high herkogamy) or from high early inbreeding depression (high abortion of seeds) (Husband and Schemske, 1996; Opedal, 2018) (Fig. 1). In contrast, predominantly selfing species or populations often exhibit reduced herkogamy and lower inbreeding depression, producing fruit and seed sets comparable to outcrossing. Consequently, the allocation to clonal reproduction may depend on the efficiency and quality of autofertility at both the plant and species level.

Species with mixed mating systems often vary in their capacity for selfing across their geographic range reflecting spatial differences in pollen limitation (Goodwillie et al., 2005). The reproductive assurance hypothesis predicts an increased capacity to self-fertilize in face of increased pollen limitation (Lloyd, 1992; Kalisz et al., 2004; Morgan and Wilson, 2005). If cloning also functions as a form of reproductive assurance, a similar increase in clonal reproduction could be expected under pollen limitation, provided that no trade-off exists between selfing and cloning (Eckert et al., 2006). Alternatively to the reproductive assurance hypothesis, pollen limitation can also increase the selective pressure for increased pollinator attraction, such as for larger petals (Emel et al., 2017).

Large-scale geographic trends in pollen limitation, such as those across latitude and altitude, have been frequently tested. If climate predicts pollinator reliability, one might expect pollination uncertainty to increase in harsher environments, such as those found closer to the pole or at higher elevations. However, our current evidence supporting this hypothesis is still unclear and dependent on a number of different factors such as life history, growth form, mating systems, pollination system, plant species richness and the extent of the geographical scale analyzed (Vamosi et al., 2006; Wirth et al., 2010; Hargreaves et al., 2015; Moeller et al., 2017; Koch et al., 2020; Dawson-Glass and Hargreaves, 2022). Overall, mating system trends at broad geographic scales remain poorly understood. Examining a species’ potential to self, to clone, and its potential to attract pollinators across its continental distribution may help reveal mechanisms maintaining mixed mating systems that include two alternative modes of reproductive assurance. Given the differential fitness repercussions associated with these two modes of reproductive assurance (Fig. 1) and their increasing relevance with the current global environmental and biodiversity changes (Rodger et al., 2021), this subject urges further investigation.

Here, we examine the relationship between autonomous selfing and cloning under experimentally induced pollinator-limited conditions in woodland strawberry (*Fragaria vesca subsp. vesca* L.) (Hilmarsson et al., 2017), a self-compatible perennial, across its distribution range. Using this approach, we test the following hypotheses: (1) clonal reproduction decreases with increasing autofertility (i.e., the capacity for autonomous self-fertilization, and (2) in harsher environments where pollinator reliability is expected to be reduced (e.g., at higher latitudes or elevations) either (i) clonal reproduction and selfing are higher, or (ii) floral display traits increase to enhance pollinator attraction. To address these hypotheses, we established a pollinator-free common garden experiment of 121 woodland strawberry genotypes, sampled from contrasting local abiotic and biotic environments across Europe (Egan *et al*. 2018; De la Cruz, Batsleer, Bonte, Diller, Hytönen, *et al*. 2025; De la Cruz, Batsleer, Bonte, Diller, Izquierdo, *et al*.2025). We quantified the capacity to autonomously self-fertilize and to clone across these genotypes with the specific aim to assess (I) the relationship between the propensity to clone and the capacity to self. In addition, we measured floral traits and conducted hand-pollinations to assess if a genotype’s capacity to self-fertilize is determined by (II) floral herkogamy, and (III) early inbreeding depression. Finally, (IV) we evaluated a genotype’s propensity to clone, its capacity to self-fertilize, and its potential for attracting pollinators (i.e. its floral display) in relation to its geographic coordinates and elevation.

## MATERIALS AND METHODS

### Study species

The woodland strawberry *Fragaria vesca* L. (see **Supplementary Section 1** for study system pictures) is a self-compatible herbaceous perennial plant (Staudt, 1989) (see **Supplementary Section 2** for hand pollination experiments). It occurs throughout the Holarctic and shows significant genetic variation in several traits, including flower traits (Egan et al., 2018, 2021). The flowers contain numerous pistils on the top of the receptacle that forms a fleshy aggregated accessory fruit (hereafter fruit) after pollination (Liston et al., 2014). The ovaries contain a single ovule and form into dry achenes located on the surface of the fruit. Each flower has about twenty stamens, arranged into two whorls of ten each. The inner whorl is subdivided in two sets of five, of different height, that alternate with each other around the pistils (see **Supplementary Section 1**, Fig. 1 D, E and F). The flowers have overlapping male and female receptivity allowing for autonomous selfing through spatial closeness between anthers and stigmas (low herkogamy). *F. vesca* plants are also able to propagate clonally through runners (**Supplementary Section 1**, Fig. 1 C). Flowers are induced by cool temperatures and short photoperiods in autumn and they emerge following spring, while the opposite conditions activate runner formation in summer (Hytönen and Elomaa, 2011).

### Plant material

Our study included 121 different genotypes/accessions of *F. vesca* collected throughout its European distribution range (Hilmarsson et al., 2017; Weber et al., 2020). A single genotype was collected from each location and sampling locations were at least a few kilometers apart from each other, except for two samples from northern Norway with a sampling distance of a about 20 meters. These genotypes were sampled between latitude 37.2° to 70.18° and longitude -23.18° to 26.07° in 2013-2014. The elevation range at which the genotypes were collected was between 0 to 2275 m.a.s.l, with a median of 60 m.a.s.l. See **Supplementary Section 3** for geographical coordinates and elevation.

### Common garden and pollinator exclusion setup

An outdoor common garden was established at the Swedish University of Agricultural Sciences in Alnarp in autumn of 2019 and data was collected during the spring-summer of 2020. The 121 genotypes were clonally replicated in the field to obtain 5 replicates (ramets/plants) per genotype. Yet due to mortality, our final set up consisted of 2 to 5 ramets (median of 3 ramets) per genotype and ramets were randomly placed across nine blocks. Our common garden consisted of a total of 363 ramets. Each ramet was individually planted in a 17 cm diameter pot with Kekkilä Professional potting substrate WR8494. Plants were watered manually as needed and covered with fleece cloth when temperatures dropped below 5 °C to protect them from frost. Pots were fertilized twice throughout the season with SW-BOUYANT RIKA S 7-1-5+MIKRO (SW Horto®). To simulate a pollinator-free environment, each plant was covered entirely with a mesh bag (45 x 60 cm organza bag Saketos Co®) to exclude pollinators. Exclusion bags were held in place and apart from the plant by plastic frames (‘Plant Supports-Series STG’ from Pöppelmann TEKU®) that fit securely on top of each pot (see **Supplementary Section** 1. Fig. 1A). They were installed just before flowering (mid-April 2020) and kept until after fruit harvest.

### Data collection

<u>Autonomous selfing</u>: to assess the capacity to autonomously self-fertilize, we randomly labeled at least one flower per ramet. Where feasible, additional flowers were labelled per ramet (one flower for 263 ramets, two for 92 ramets, and three for eight ramets). For each fruit, we recorded its position within the inflorescence to account for within-inflorescence variation in downstream analyses (see below) (Guitián and Navarro, 1996). Fruits were harvested and kept in separate vials in a freezer until processed in the lab. We then counted the achenes and categorized them under a dissecting scope as fertilized or not, based on their shape and size. Achenes that were visibly bigger and swollen were considered as fertilized (MacInnis and Forrest, 2017). The proportion of fertilized achenes of these autonomous fruits was used as a proxy for a genotype’s capacity to self-fertilize. Finally, we also measured the size of the fruits (fruit volume = width_1_ * width_2_ * height). Fruit measurements were taken with a digital caliper with an accuracy of 0.01 mm.

#### Cloning

To assess a genotype’s propensity to clone, we counted the total number of runners produced by each ramet from the onset of flowering to the end of the summer (see below for more details).

#### Flowering

We counted the total number of flowers per ramet. As for runner production, flower number was counted from the onset of flowering to the end of the summer. Both flower and runner numbers were measured three times during the flowering season: early, peak, and late. Early-season measurements took place between 28^th^ April and 9^th^ of May, mid-season between 25^th^ and 29^th^ of May, and late season between 17^th^ and 28^th^ of July. We used the cumulative value both for flower and runner numbers in our analyses.

#### Floral herkogamy

To evaluate whether herkogamy facilitates autonomous self-fertilization, we measured the distance between anthers and pistils (Toräng et al., 2017; Opedal, 2018). We measured this distance both in the horizontal (lateral herkogamy) and vertical plane (vertical herkogamy) (see **Supplementary Section** 1). Flower traits were measured during the peak flowering time (8^th^ May – 4^th^ June). Wherever possible, we measured two flowers per ramet and, again, took note of the position of the flower within the inflorescence. Despite these efforts, the median number of flowers measured per ramet was equal to one. We measured a total of 311 flowers across a total of 363 ramets. Measurements were taken only on fresh flowers with fully stretched out petals and stamens. We used a digital caliper (0.01 mm) and measured every trait twice to account for measurement error and later averaged. To improve the precision, floral measurements were destructive. Therefore, floral and fruit measurements were not taken on the same flower.

#### Floral display

To assess the potential of a genotype to attract pollinators, we measured petal size and the inflorescence exsertion (i.e. the extension of the inflorescence above the leaf canopy), in addition to counting the total number of flowers per ramet (see above). Petal size was measured at the same instance and following the same criteria described above for lateral and vertical herkogamy. Inflorescence exsertion was measured on the highest inflorescence at the end of the flowering season.

#### Plant size

We calculated plant volume as a proxy for plant size (used as a covariate to account for variation in biomass) as the product of the two perpendicular width measurements and the height of the leaf rosettes. Measurements were taken during the late season (17^th^ – 31^st^ of July).

#### Inbreeding depression

To assess the degree to which autonomous selfing is limited by early inbreeding depression (i.e. embryo abortion), we calculated the relative performance (RP) for each genotype (see below) with the results obtained from outcross and assisted self-cross hand pollination treatments. In contrast to the traditional estimation of inbreeding depression, the advantage of utilizing the RP index is its symmetrical distribution and that it is bounded by -1 and 1 (Ågren and Schemske, 1993; Dudash et al., 1997). Positive values indicate greater inbreeding depression (i.e. reduced fitness for selfing when compared to outcrossing). Following Ågren and Schemske (1993), we calculated the relative performance (RP) as:

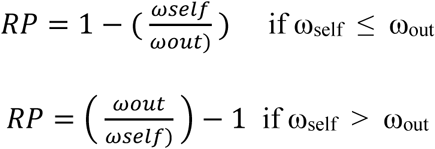

Where ω_self_ and ω_out_ is the fitness measured for selfing fruits and out-cross fruits, respectively. Outcrossing consisted of hand pollinations with pollen collected from a different genotype and selfing consisted of hand pollinations with pollen collected from the same flower or flowers from the same ramet. Fitness differences between assisted selfing and outcrossing were assessed by the proportion of fertilized achenes of each treated fruit. We averaged across fruits whenever the ramet had more than one fruit per treatment. Due to the limited flower availability, we were able to calculate this index only for a subset of 34 genotypes (84 ramets), for which we had data for both within-individual (ramet) selfing and out-cross hand pollinations (see **Supplementary Section 4**, for genotype list and description). We only included genotypes for which we had RP values for at least two replicates (two ramets) per genotype. Finally, to be able to compare the overall degree of inbreeding depression in *F. vesca* compared to other reported systems, we also calculated the traditional estimate of inbreeding depression (δ) as follows:

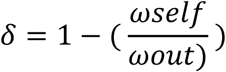

## STATISTICAL ANALYSIS

### Statistical framework and preliminary analyses

All statistical analyses and graphs were done with JMP Student Edition 18.2 (JMP®, n.d.). Maps were generated in R (R CoreTeam, 2021) with ggplot2 (Wickham., 2016) and terra packages (Hijmans, 2025). Figure panels were edited with Adobe Illustrator 2020. All analyses consisted of mixed models with genotype and block as a random term. Whenever more than one measurement was recorded per ramet, values were averaged at the ramet level. All explanatory variables were standardized to a mean of zero and standard deviation of one. We examined the residual distributions of all models to verify that model assumptions were met (normality of residuals through the QQ plot and homoscedasticity through the residual by predicted plot). All models were fitted as linear mixed models (LMM), except when the response variable consisted of count data (runner and flower number). In those cases, we used generalized linear mixed models (GLMM) with negative binomial distribution to account for overdispersion. When modelled as a response variable, proportion of fertilized achenes was logit-transformed to satisfy linear mixed model assumptions.

Since *F. vesca* inflorescences consist of primary, secondary, and tertiary flowers that visibly vary in size, we first ran exploratory linear mixed models to assess the effect of inflorescence position on floral traits before conducting the main analyses addressing our research questions. We found no significant effect of inflorescence position on lateral and vertical herkogamy (see **Supplementary Section** 5, Table 1) and hence excluded flower position from any further model concerning lateral or vertical herkogamy as fixed factors. Petal size, however, was significantly smaller in tertiary flowers than in primary or secondary flowers. To account for this, all further models that included petal size as the response variable were done with the conditional predicted values obtained from the model described in the **Supplementary Section** 5, Table 1A and D. Similarly, we also checked for the effect of inflorescence position for traits measured on autonomous fruits (i.e. fruit size and the proportion of fertilized achenes). Autonomous fruit traits did not vary in relation to inflorescence position and hence was excluded from any further analyses (see **Supplementary Section** 5, Table 2). Finally, we tested for multicolinearity among explanatory variables by running Pearson correlations and found no evidence of multicollinearity (i.e. all r < 0.7) (see **Supplementary Section** 5, Table 3) (Dormann et al., 2013).

**Table 1.**
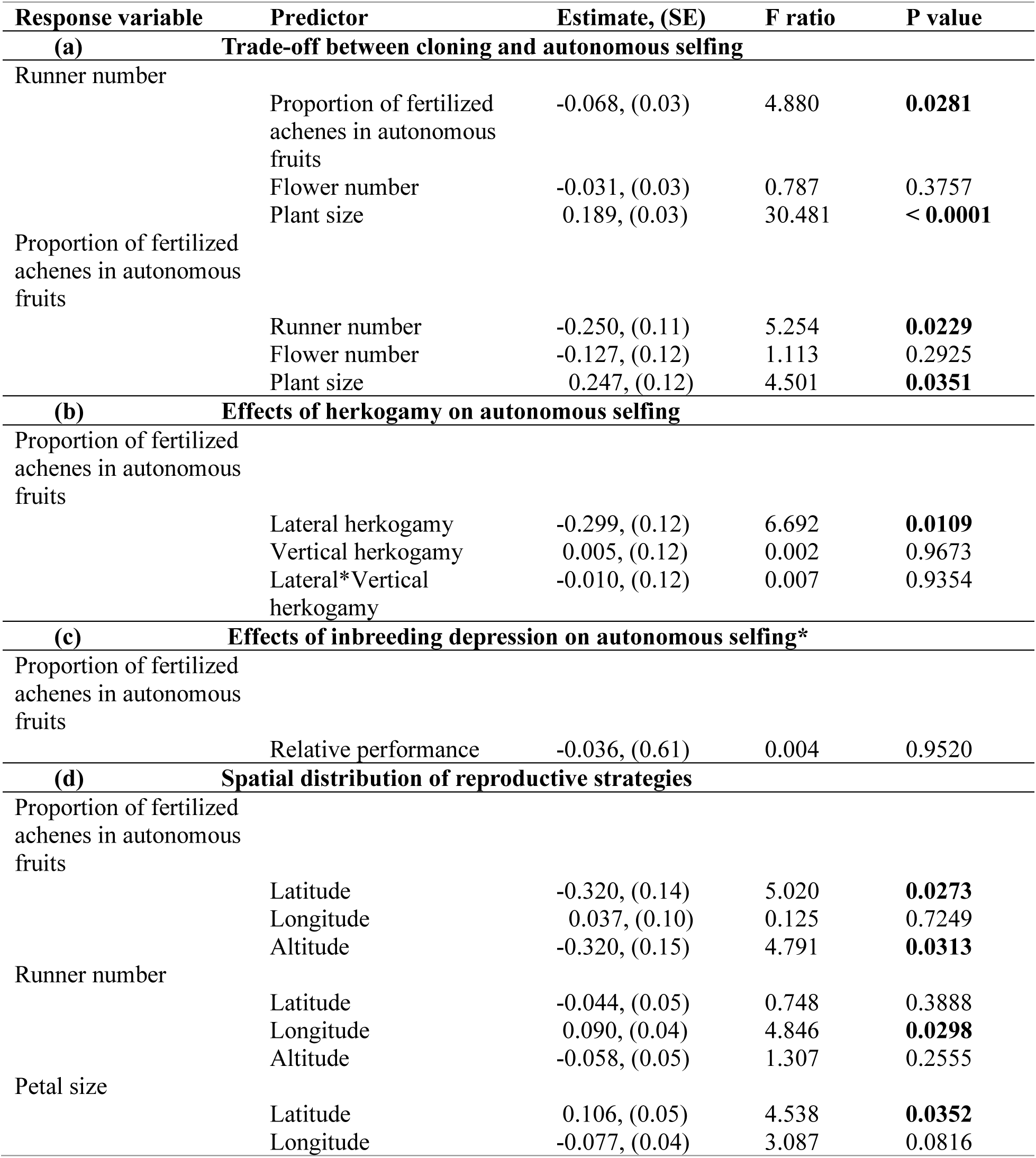
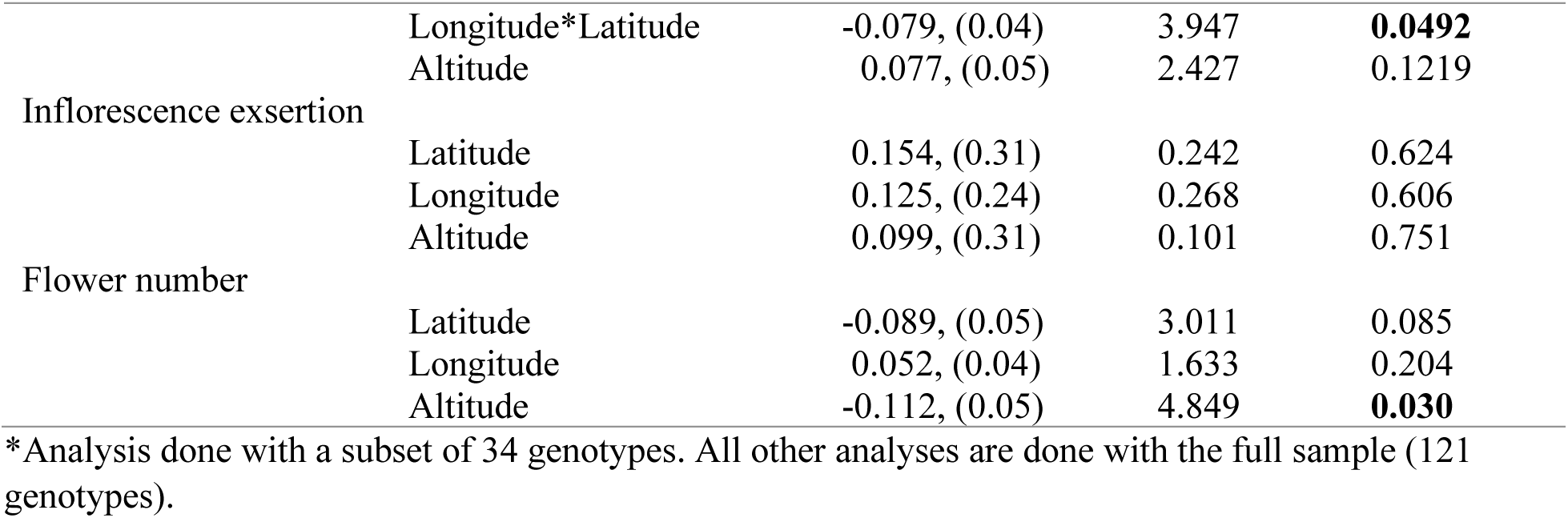
Mixed-models testing the **(a)** trade-off between cloning and autonomous selfing, **(b)** effects of herkogamy on autonomous selfing**, (c)** effects of inbreeding depression on autonomous selfing and **(d)** spatial distribution of reproductive strategies. We refer to runner number as a proxy for the propensity to clone, the proportion of fertilized achenes on autonomous fruits as a proxy for the capacity to autonomously self-fertilize and traits related to floral display (petal size, inflorescence exsertion and flower number) as proxies for the attractiveness to pollinators.

### Reproductive assurance allocation: i) selfing vs. cloning

To test for a trade-off between the capacity to self-fertilize and the propensity to clone across genotypes, we performed a generalized linear mixed model with runner number as the response variable and the proportion of autonomous achenes as the explanatory variable. We included plant size as a covariate, to account for differences among plant individuals (ramets) in resource availability or allocation to other organs. Flower number was also added as a covariate in the model, to control for the possibility of resource limitation by previously invested resources into producing flowers. See **Supplementary Section** 5, Table 4 A for model details. We also ran an inverted linear mixed model (with the proportion of fertilized achenes as the response variable and runner number as the explanatory variable) to corroborate if the trade-off occurs both ways. See **Supplementary Section** 5, Table 5 A for model details. Finally, to gain further insight into the potential trade-off between selfing and cloning, we tested for a correlation between fruit size and the proportion of fertilized achenes using a Pearson correlation, as a positive relationship would indicate increased resource investment in fruit development with higher achene set.

### Capacity to self-fertilize: i) floral traits

To assess whether floral traits attributed to increased selfing indeed facilitate selfing in *F. vesca* we ran a linear mixed model with the proportion of fertilized achenes as the response variable and lateral and vertical herkogamy as explanatory variables. See **Supplementary Section** 5, Table 6 A for model details.

### Capacity to self-fertilize: ii) inbreeding depression

To evaluate whether inbreeding depression influences the capacity to self-fertilize across genotypes, we performed a linear mixed model with the relative performance (calculated based on the proportion of fertilized achenes from assisted selfing and outcross hand pollinations) as predictor and the proportion of autonomous fertilized achenes as response variable. See **Supplementary Section** 5, Table 7 A for model details.

### Large scale geographical trends

To examine whether the capacity to autonomously self, the propensity to clone and the potential to attract pollinators within *F. vesca* is driven by its geographical coordinates or elevation, we performed five mixed models. The predictors in all models consisted of the standardized values of latitude, longitude and altitude. The interaction term between latitude and longitude was included if significant. We did not incorporate an interaction term with altitude in the model, due to lack of replication with varying altitudes among our genotype samples (see section: *‘Plant material’*). The response variables included the proportion of autonomous fertilized achenes, the number of runners, petal size, inflorescence exsertion and flower number. The last three were considered as descriptors of floral display and hence as proxies for the potential to attract pollinators. See **Supplementary Section** 5, Table 8 A, 9 A and 10 A for model details.

## RESULTS

### Reproductive assurance allocation: i) selfing vs. cloning

After accounting for plant size and flower number, genotypes more capable of self-fertilizing were less prone to cloning (Table 1; see **Supplementary Section** 5, Table 4 B for full model output). Specifically, higher proportions of fertilized achenes were associated with reduced number of runners (approximately 7% decrease per standard deviation increase in achene set; Fig. 2. Panel I). Inverting the model, with the proportion of fertilized achenes as the response variable and runner number as the explanatory variable resulted in a similar negative relationship (Table 1; see **Supplementary Section** 5, Table 5 B for full model output). The total number of flowers per ramet was not significantly associated with runner numbers or achene set of self-fertilized fruits. Fruits with a higher achene set were in general larger in size (Pearson correlation r = 0.51; Fig. 2. Panel II; **Supplementary Section** 5, Table 11).

**Fig. 2.**
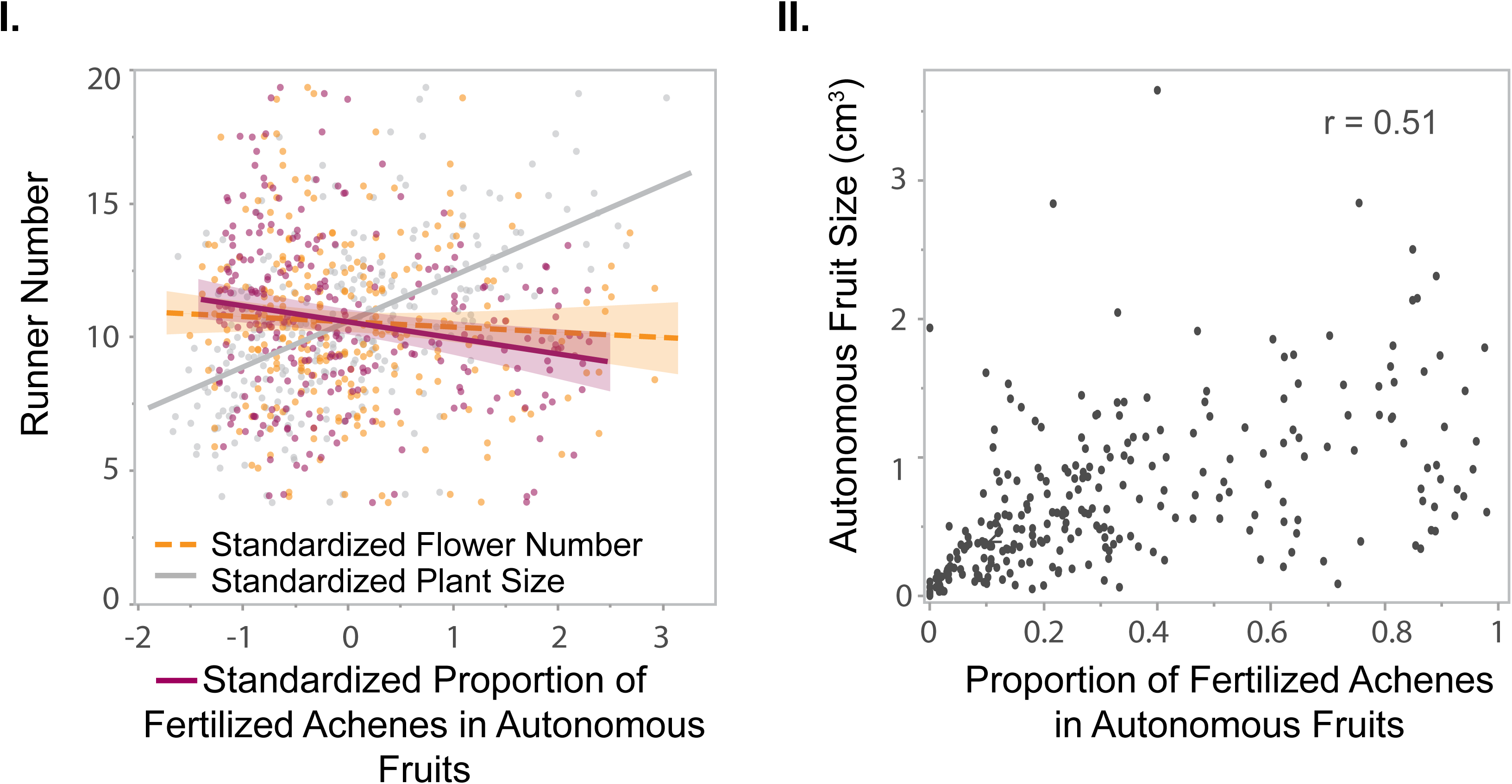
*Fragaria vesca* genotypes that are better at selfing are worse at cloning (Panel I), likely due to increased resources required to produce large fruits with high achene set (Panel II). In panel I and II each dot represents a ramet belonging to 121 *F. vesca* genotypes and measured under a pollinator-free common garden. Panel I: the capacity to autonomously selfing is quantified as the proportion of fertilized achenes per autonomous fruit and the propensity to clone is quantified as the number of runners produced per plant. Lines show conditional predictions from a negative binomial GLMM with standardized fixed effects. Solid lines are statistically significant and dashed lines are non-significant (see Table 1 and **Supplementary Section** 5, Table 4). The x-axis is standardized. Mean and standard deviation of predictor variables in original units: 4052 ± 2242 cm^3^ for plant size, 25 ± 17 for number of flowers and 0.34 ± 0.28 for the proportion of fertilized achenes. Panel II: pearson correlation between autonomous achene set and fruit size. See **Supplementary Section** 5, Table 11.

### Capacity to self-fertilize: i) floral traits

Flowers with reduced lateral herkogamy produced autonomous fruits with a higher proportion of fertilized achenes (Table 1 and Fig. 3. Panel I). Vertical herkogamy, however, did not influence the proportion of fertilized achenes in autonomous fruits (Table 1 and Fig. 3. Panel I). See **Supplementary Section** 5, Table 6 B for full model output.

**Fig. 3.**
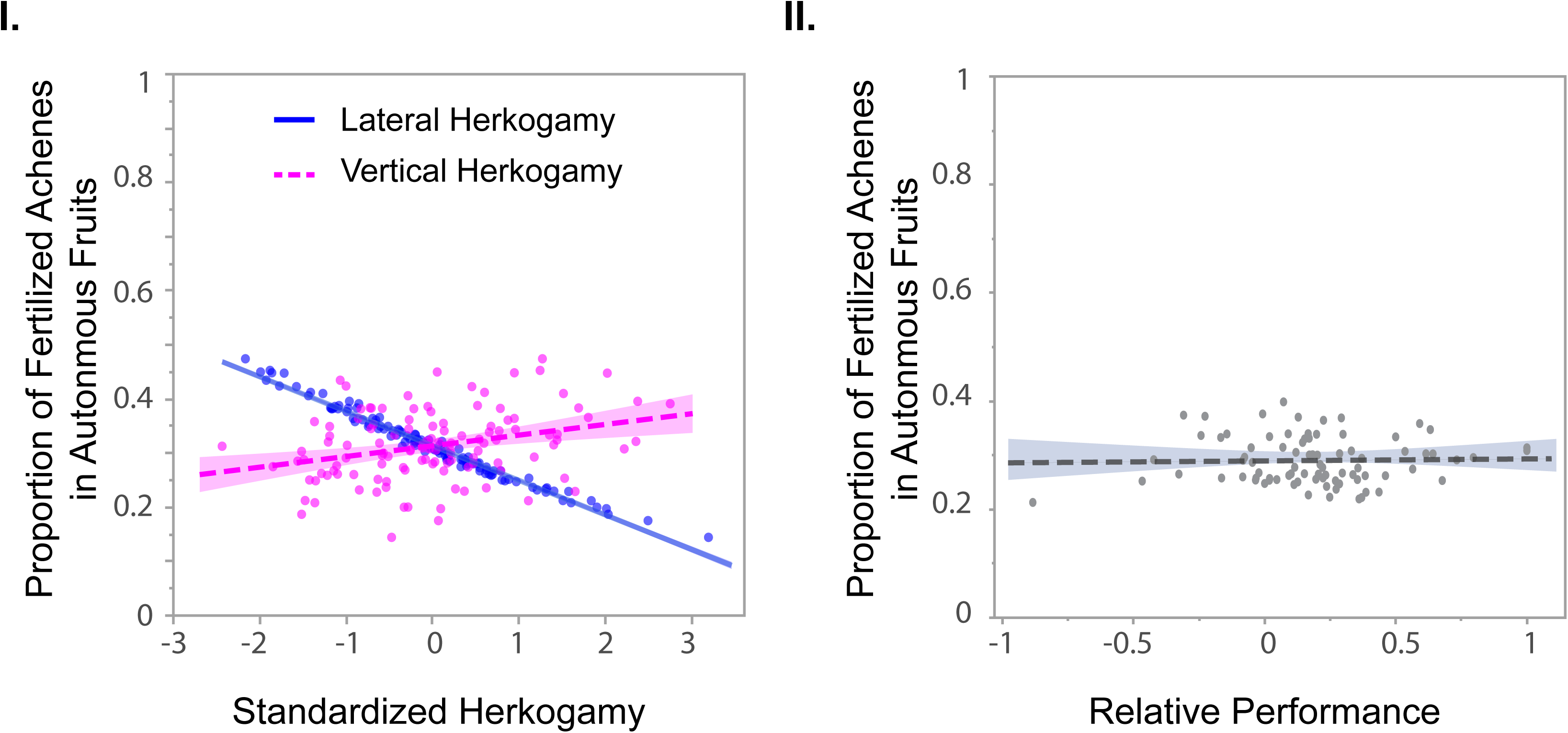
The capacity to self-fertilize is defined by lateral herkogamy (Panel I), but not by inbreeding depression (Panel II) in *Fragaria vesca*. Panel I: The capacity to self-fertilize is quantified as the proportion of fertilized achenes in autonomous fruits. Y axis are back transformed conditional predicted marginal means. Each dot corresponds to a ramet corresponding to 121 genotypes. Solid lines are statistically significant and dashed lines non-significant. See Table 1 and **Supplementary Section** 5, Table 6. The x-axis of panels is standardized. Mean and standard deviation of predictor variables in original units: 0.62 ± 0.33 cm in lateral herkogamy, and 0.06 ± 0.49 cm in vertical herkogamy. Panel II: Inbreeding depression was quantified as the relative performance (RP) across a subsample of 34 genotypes. Each dot corresponds to a ramet. Positive values of RP indicate higher inbreeding depression (see methods). See **Supplementary Section** 5, Table 7.

### Capacity to self-fertilize: ii) inbreeding depression

Across the subsample of 34 genotypes, mean inbreeding depression was 0.18 ± 0.04 (range −0.87 to 1.00), indicating that assisted selfing resulted in 18% lower number of fertilized achenes relative to hand-outcrossing. Nevertheless, inbreeding depression (quantified as the relative performance) did not influence the capacity to autonomously self-fertilize (Table 1 and Fig. 3, Panel II). See **Supplementary Section** 5, Table 7 B for full model output.

### Large scale geographical trends: i) reproductive assurance

The capacity to autonomously self-fertilize decreased with increasing latitude or altitude, whereas longitude had no effect (Table 1 and Fig. 4. Panel I and II). See **Supplementary Section** 5, Table 8 B for full model output. The propensity to clone, on the other hand, was associated only with longitude, increasing towards the east (Table 1 and Fig. 4. Panel III and IV). See **Supplementary Section** 5, Table 9 B for full model output.

**Fig. 4.**
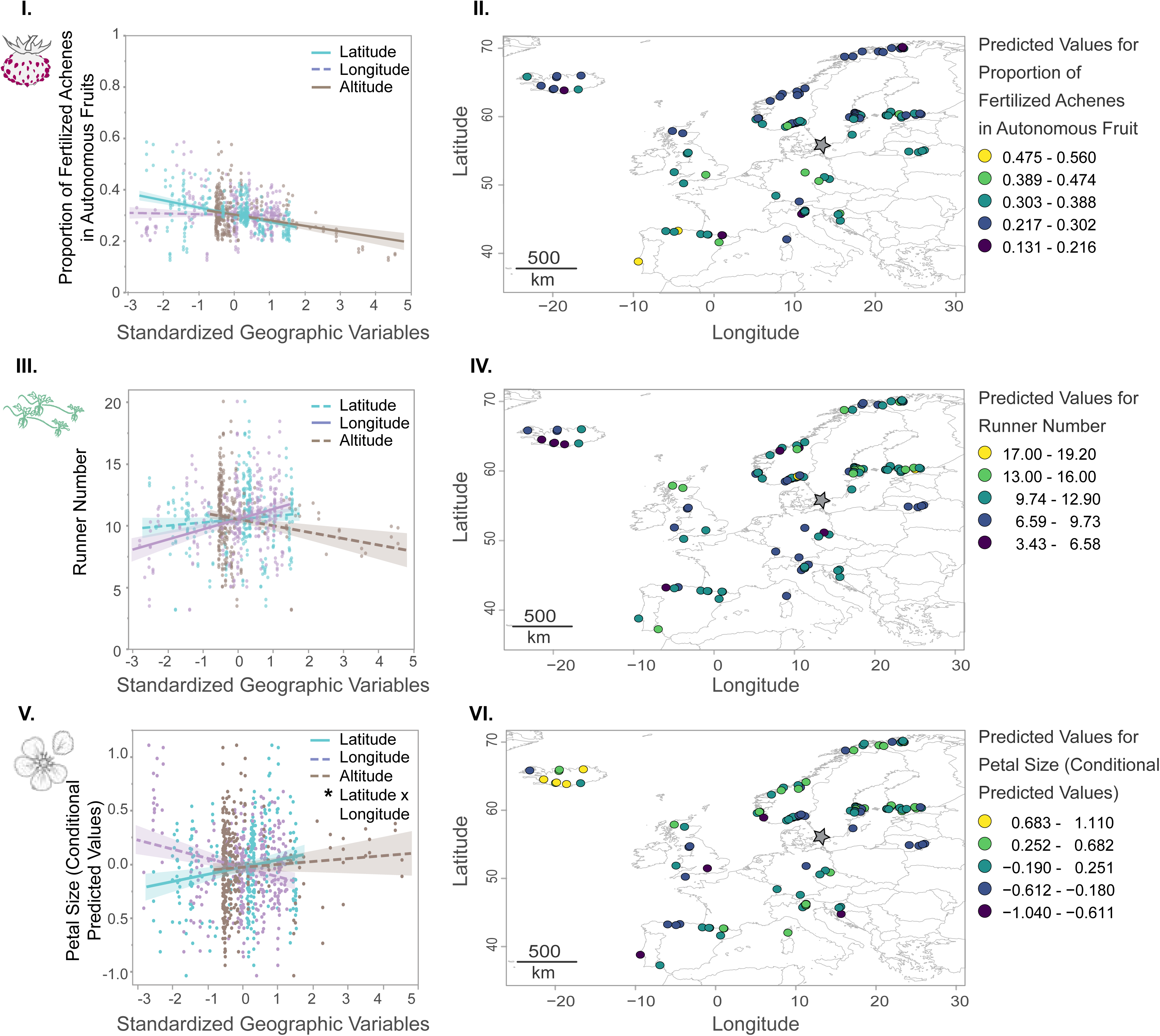
Geographic patterns in reproductive traits of *Fragaria vesca* under a pollinator-free common garden. Genotypes sampled at lower latitudes and altitudes showed a higher capacity for autonomous self-fertilization (Panels I–II), whereas genotypes sampled further east exhibited a greater propensity for clonal reproduction (Panels III–IV). Petal size was largest in genotypes originating from the northwest and smallest from the southwest (Panels V–VI). Autonomous selfing was quantified as the proportion of fertilized achenes per autonomous fruit, and clonal propensity as the number of runners produced per plant. Petal size was used as a proxy for pollinator attraction. Panels I, III, and V show regression plots (Table 1 and **Supplementary Section 5**, Tables 8–10), with each point representing a ramet from 121 genotypes. Solid lines indicate statistically significant effects, whereas dashed lines indicate non-significant effects. Predictor variables are standardized (mean ± SD in original units: latitude 57.6 ± 8.12°, longitude 10.6 ± 12.14°, altitude 243 ± 448 m a.s.l.). The y-axis in Panel I shows back-transformed conditional predicted marginal means, whereas Panel V shows the conditional predicted values from the model correcting for flower position (see **Supplementary Section 5**, Table 1). Panels II, IV, and VI show predicted values mapped across Europe. Circles indicate the sampling locations of genotypes (n = 121), and the grey star marks the location of the common garden.

### Large scale geographical trends: ii) floral display (proxies for pollinator attraction)

Overall, genotypes sampled towards the north had larger petals. The effect of longitude on petal size depended on latitude (latitude x longitude interaction) (Table 1 and Fig. 4. Panel V and VI.). Among the northern genotypes, petal size increased strongly towards the west, while in the southern genotypes a slight increase of petal size was found towards the east (**Supplementary Section** 5, Fig. 1). Consequently, petal size was greatest in the northwest and smallest in the southwest, reflecting the stronger westward increase at higher latitudes compared to the weaker eastward increase at lower latitudes. See **Supplementary Section** 5, Table 10 B for full model output. Genotypes sampled at lower elevations produced more flowers, however, inflorescence exsertion did not vary with latitude, longitude or altitude (Table 1; see **Supplementary Section** 5, Table 10 C and D for full model output).

## DISCUSSION

### Reproductive assurance allocation: selfing vs. cloning

Overall, the genotypes that produced good quality autonomous fruits (i.e. fruits with high achene set), produced lower numbers of runners. It is possible that resource competition may have played a role in this trade-off given that higher achene set resulted in larger fruits (Fig. 2. Panel II). Similar trade-offs between runners and fruiting take importance in commercial strawberry production (Khan et al., 2023). Notably, we detected a trade-off between sexual reproduction and cloning even though fruits produced through autonomous selfing are significantly smaller than fruits produced through outcrossing (**Supplementary Section 2**). Moreover, because our estimates are based on only one to two fruits per individual (ramet), rather than total fruit production and corresponding achene set, they likely represent a conservative assessment. Despite this, a clear trade-off was still evident. Finally, the negative relationship between selfing quality and cloning was not mirrored by a corresponding trade-off between flower and runner production, nor was selfing capacity related to flower number. Overall, our results suggested that fruit development rather than flower production explained the observed trade-off between the two modes of reproductive assurance.

Alternative to resource competition, reproductive trade-offs are often also explained by a competition of meristems (Geber et al., 1992). In *F. vesca*, the inflorescence and flower meristems are initiated in the autumn, before pollinator limitation is experienced in the spring when they flower (Hytönen and Kurokura, 2020). Runners on the other hand, are initiated from axillary meristems during the spring and summer time (Hytönen and Kurokura, 2020; Andres et al., 2021). Since inflorescences and runners do not originate from the same meristems in *F. vesca*, we rule out meristem competition as an explanation for the observed trade-off. Furthermore, even if one meristem type could developmentally arrest another when active (Geber et al., 1992), this would manifest as a trade-off between flowers and runners, rather than between the quality of fruits and runners.

### Capacity to self-fertilize

As expected, we found that genotypes had a greater capacity to self-fertilize (increased proportion of fertilized achenes) when anthers were in close lateral proximity to the stigmas. Interestingly, vertical herkogamy did not contribute to the ability to self despite this being true for other systems (Brys and Jacquemyn, 2012; Opedal, 2018; Jiménez-López et al., 2020) (Fig. 3. Panel I) and emphasizes the importance to look at lateral herkogamy in addition to vertical herkogamy, which is often overlooked (but see Jiménez-López et al., 2020). In a competing selfing system such as in *F. vesca*, where outcrossing and selfing may occur simultaneously, one expects conflicting selection between traits that favor selfing vs. outcrossing (Lloyd, 1992). Since the genotypes included in this study were not phenotyped under natural open pollination conditions, we cannot state that the opposite trait values (i.e. greater lateral anther/stigma separation) indeed produce higher outcrossing achene set.

Finally, the capacity to self-fertilize was not limited by the level of inbreeding depression, likely due to the overall low inbreeding depression found across our study system.

### Reproductive modes across large scale spatial gradients

Following the reproductive assurance hypothesis, one would expect genotypes to be better at cloning and selfing in regions where we anticipate less reliable pollination service (i.e. higher pollen limitation) due to harsh weather, such as towards the arctic or at high elevations.

Contrary to this hypothesis, the ability to self-fertilize was reduced with increasing latitude and altitude. These findings either suggest that there is no pollen limitation towards harsher climates, or that pollen limitation is not selecting for increased reproductive assurance.

However, a recent population genomic study on *F. vesca* - that includes about 60% of the genotypes included in this study - found increased inbreeding coefficient (F_IS_) and reduced effective population size (N_E_) with increasing latitude (Toivainen et al., 2026). This suggests that at more northern latitudes, populations are likely small and fragmented with reduced mate diversity. Since, pollen limitation can result due to low pollen quantity (pollinator availability) or low quality (low mate availability) (Aizen and Harder, 2007), it is plausible that the more northern populations are indeed more pollen limited. If pollen limitation is in fact stronger in northern populations, this lends support to the latter explanation, that pollen limitation is not selecting for increased reproductive assurance via selfing.

Genotypes had larger petals at higher latitudes, likely as a result of increased pollinator-mediated selection for increased floral attraction and hence increased outcrossing. This is in attune with other findings that found increased pollen production at higher latitudes (Cunha et al., 2022) and increased stigma receptivity (floral longevity) in alpine plants (Bingham and Orthner, 1998) increasing the pollination efficiency and reducing pollinator limitation due to pollination unpredictability in these areas (Bingham and Orthner, 1998; Koch et al., 2020). Since outcrossing and selfing occur simultaneously and hence compete in *F. vesca*, our results suggest that the likely pollen limitation experienced by *F. vesca* at higher latitudes (either due to reduced mate diversity or increased pollinator uncertainty) is enhancing selective pressures in favor of outcrossing (larger petals) rather than increased reproductive assurance through autonomous selfing.

The propensity to clone, however, did not vary with latitude or altitude. Instead, it increased with genotypes sampled towards the east. We have no prior expectations regarding pollen limitation across a longitudinal scale in terms of pollinator uncertainty. However, we know that the eastern populations of *F. vesca* have much higher effective population sizes, e.g., in Baltic countries or southeastern Europe, than western populations (Toivainen et al., 2026), implying lower mate availability and hence also possibly higher pollen limitation towards the west. Interestingly, at least at higher latitudes, we found increased potential to attract pollinators (larger petals) also toward the west. Following the same logic as above, we propose that this higher pollen limitation towards the west may be enhancing selecting pressures in favor of pollinator attraction through larger flowers rather than increased reproductive assurance through cloning (Cisternas-Fuentes and Koski, 2024). Alternatively, these longitudinal patterns may reflect differentiation among genetic clusters in western and eastern Europe resulting from past glacial refugia (Toivainen et al., 2026) or from variation in abiotic conditions (Zhang et al., 2023; Chen et al., 2025).

We also found increased flower production in genotypes sampled at lower elevations. Overall, we expected floral display traits to show similar spatial patterns, reflecting a shared strategy to enhance pollinator attraction. Although most spatial effects were not statistically significant, the slopes for flower number and petal size were consistently in opposite directions (Table 1), hinting at a potential trade-off between flower size and number (Sargent et al., 2007; Goodwillie et al., 2010). However, patterns involving altitude should be interpreted with caution as our sampling was strongly skewed towards lower elevations. Some caution is also warranted when interpreting our longitudinal gradient, which was well represented at higher latitudes but less so in southern Europe.

A caveat in these interpretations is that we do not provide evidence for the cause of these reproductive gradients across space. In particular, we do not provide evidence that larger petals indeed attract more pollinators in *F. vesca*. Nevertheless, assessing the spatial dimension to our analyses revealed a relevant point to the understanding of the role of selfing and cloning as two alternative modes of reproductive assurance. If cloning and selfing served as two redundant modes of reproductive assurance, all genotypes would be able to clone and self equally well. However, the reproductive trade-off detected in this study between cloning and selfing informed us that some genotypes are better at selfing, whereas others are better at cloning. In other words, we identified that a genotype’s propensity to clone is conditioned by its ability to self. Within this framework, one would expect selfing and cloning to vary inversely across the same gradient (e.g. latitude). However, here we find that this trade-off in modes of reproductive assurance is split across the latitude and longitude. Selfing is more dominant at the latitudinal (and altitudinal) gradient and cloning more towards the longitudinal gradient.

### Additional evolutionary nuisance on cloning

In this study we discuss some of the drivers that may shape the mating system in a species that can outcross, self and clone depending on whether pollinators are present or absent. However, the scenario may be more nuanced as clonality can contribute additional disadvantages not depicted in Fig.1. For instance, clonality can reduce the effective population size and hence increase the genetic load of a population or reduce the rate at which lethal alleles are purged from the population (Marriage and Kelly, 2009). Additionally, clonality can also increase inbreeding depression levels through increased life-span and or somatic mutations (Muirhead and Lande, 1997). Therefore, increased clonality could interfere with the rate at which lethal alleles are purged through selfing and hence also reduce the rate at which a population can increase its capacity to self-fertilize (Marriage and Kelly, 2009). Moreover, cloning can also ultimately increase sexual reproductive capacity of a genotype, as every clone will ultimately also produce flowers that will either self or outcross (Vallejo-Marín et al., 2010). More empirical evidence is needed to understand these broader impacts of cloning in a mixed mating system.

## Conclusion

Altogether, when assessed at the species level, we found a trade-off in the woodland strawberry’s expression of autonomous selfing versus cloning when pollinators are absent. Genotypes with traits favoring selfing (e.g. reduced lateral herkogamy) produce autonomous fruits with higher proportion of fertilized achenes. Since berries are larger with a higher achene set, it is likely that these bigger fruits may compete for resources for runner production. Important in the above narrative and keeping in mind that we found that the trade-off is not defined by the amount of flowering, we show that it is ultimately the ability to self that positions a genotype within the selfing/cloning gradient trade-off. Interestingly, when the reproductive allocation was analyzed spatially at a continental scale, the modes of reproductive assurance split across latitude and longitude. The capacity to self-fertilize increased at lower altitudes and towards the South of Europe, while the propensity to clone increased towards the East of Europe, compensating perhaps for the reduced attractiveness to pollinators (smaller petals) of the genotypes sampled from these respective areas.

## ACKNOWLEDGEMENTS

We thank Maja Brus-Szkalej and Valeria Rossi for help collecting data in the field, Adam Flöhr for statistical advice on an early version of this article, Guillermo Rehermann for valuable input during the preparation of this manuscript, Cajsa Lithell for illustrative advice on Fig. 1, and Åsa Lankinen and four anonymous reviewers for helpful comments on an early version of this article.

## AUTHOR CONTRIBUTIONS

Carolina Diller Johan A. Stenberg, and Paul A. Egan conceived the project; Timo Hytönen. provided the plant collection; Johan A. Stenberg, Paul A. Egan replicated the plants onsite; Carolina Diller collected the data, wrote the first full draft and created the figures; Carolina Diller and Ivan M. De la Cruz performed the statistical analyses. All authors contributed to the interpretation of the data and critically revised the text.

## DATA AVAILABILITY

Data will be archived in a publicly available data repository upon acceptance.

## AI ASSISTANCE ACKNOWLEDGEMENT

We used ChatGPT image generator 4o to generate the hoverfly illustration (insert in Fig. 1) and the flower illustration (insert in Fig. 4.V).

## SUPPLEMENTARY INFORMATION

S.I. Section 1. Study system and flower details.

S.I. Section 2. Breeding system of *Fragaria vesca*.

S.I. Section 3. Geographical coordinates of genotype samples.

S.I. Section 4. Subset of genotypes included in inbreeding depression analysis.

S.I. Section 5. Summary statistics.

## Supplementary Section 1. Study system and flower details

**Supplementary Information Section 1. Fig. 1.**
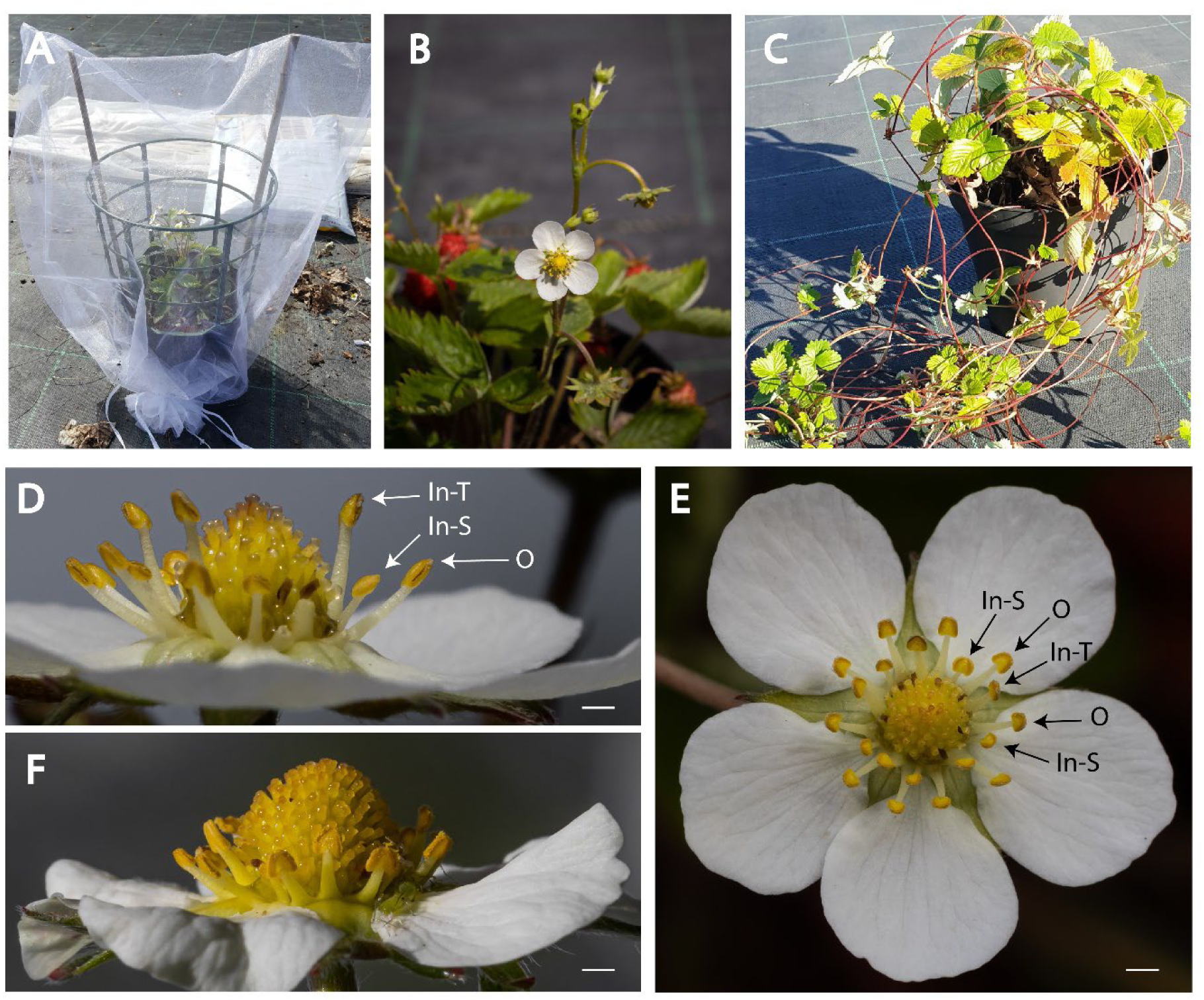
*Fragaria vesca*. Pollinator exclusion set up (A), flowering and fruiting plant (B), plant with runners (C), and flower (D, E and F). In panel (D), stamens were removed on the right side of this flower to show the three stamen types in a *F. vesca* flower: the outer whorl (O), the inner whorl, tall stamen (In-T) and the inner whorl, short stamen (In-S). Measurements for herkogamy were done on the tallest inner stamen (In-T), at the base of the anther. (F) Example of approach herkogamy (i.e. pistils higher than anthers). Scale bar in (D-F) equals 1mm. *Photo credits* for (B, D - F) Leif Elison.

## Supplementary Section 2. Breeding system of *Fragaria vesca*

### Methods

#### Hand pollination treatments

To formally describe the breeding system of *Fragaria vesca,* we performed four pollination treatments on each ramet out of 121 genotypes: outcross, self, autonomous and emasculation. Outcrossing consisted of performing hand pollinations with pollen collected from a different genotype; selfing consisted of hand pollinations with pollen collected from the same flower or flowers from the same ramet; autonomous consisted of unmanipulated flowers; and finally emasculation consisted of flowers on which we removed undehisced anthers at bud stage with forceps and left unmanipulated after that. All flowers were excluded from pollinators prior and after the treatment application.

On occasion, some pollination treatment were performed twice on a given ramet. We recorded the flower position of each hand pollination. Due to limited availability of flowers we were not able to perform all four pollination treatments for every ramet, resulting in an unbalanced sample size for all treatments. In total, we performed 363 autonomous, 361 emasculated, 313 self and 179 outcross hand pollinations.

All treated flowers were checked on a continuous basis for fruit formation. Fruits were harvested as they matured and kept in separate vials in a freezer until processed in the lab. Each fruit was then measured with a digital caliper (two measurements of width and one of length). We calculated fruit size by multiplying the three fruit measurements. Finally, we counted all the achenes on each harvested fruit and categorized the achenes under a dissecting scope as fertilized or not, based on their shape and size. In sum, for each hand pollination, we obtained values for fruit set, fruit size and the proportion of fertilized achenes.

#### Achene germination

To corroborate that our categorization of fertilized and non-fertilized achenes was correct, we set up a germination test. We randomly selected subsample of 12 fruits of each hand pollination treatment (harvested all from the same block). Each fruit belonged to a different genotype. From each fruit, we randomly selected up to 20 fertilized or non-fertilized achenes and put them in a petri dish with a moist filter paper. Similar to Chambers et al.’s (2013) protocol, we sterilized achenes in 70% ethanol for 5 min, cleansed them in 1% sodium hypochlorite for 20 min, and finally rinsed them six times in sterile, distilled water before placing them in their respective petri dishes. The petri dishes were placed in a growth chamber at 25 °C, under an 18 hr day / 6 hr night fluorescent lighting set up. Achenes were checked on a weekly basis for a period of three weeks. We considered a germinated achene only if the cotyledons emerged.

### Statistical Analysis

To assess the breeding system of our population in terms of quantity and quality of fruit, we ran three models. The first model included fruit set as the response variable, the second model fruit size and the third model the proportion of fertilized achenes (see Supplementary Section 2. Table 1. A for model description). We ran a generalized linear mixed model with a binomial distribution for fruit set, and a linear mixed model for both fruit size (log transformed) and the proportion of fertilized achenes (logit transformed). Values were averaged at the ramet level before running all analyses. The explanatory variables in the models consisted of hand pollination treatments (four levels: autonomous, emasculation, self cross and out cross) and plant size (standardized). We also included fruit position within the inflorescence (i.e. primary, secondary or tertiary) as a covariate in the models for fruit size and for the proportion of fertilized achenes. We did not include interactions across the explanatory variables, as both plant size and fruit position were only included as covariates for the purpose of these analyses. Finally, we performed a post hoc Tukey test for all three models.

### Results

#### Hand pollinations

Fruit set, fruit size and the proportion of fertilized achenes differed significantly across hand pollination treatments (F_(3, 1211)_ = 82.86, P < 0.0001, F_(3, 823)_ = 86.67, P < 0.0001 and F_(3, 739)_ = 201.64 and P < 0.0001; respectively). Overall, ‘self-cross’ and ‘out-cross’ hand pollinations were better at setting fruit (mean [LSE, USE]: 0.95 [0.94, 0.96] and 0.97 [0.96, 0.98] respectively), set the largest fruit (mean [LSE, USE]: 72.7 cm^3^[60.8, 86.8] and 93.6 cm^3^ [77.7, 112.8] respectively), and had the highest proportion of fertilized achenes (mean [LSE, USE]: 0.65 [0.6, 0.7] and 0.83 [0.8, 0.86] respectively). Compared to flowers that were hand pollinated with out-cross and self-cross pollen, flowers under the ‘autonomous self-fertilization’ treatment produced 10% less fruit (mean [LSE, USE]: 0.8 [0.77, 0.83]; Tukey: P < 0.0001), fruits a third to half the smaller in size (mean [LSE, USE]: 50.6 cm^3^ [42.5, 60.2]; Tukey: P < 0.0001) and with at least half the proportion of fertilized achenes (mean [LSE, USE]: 0.32 [0.28, 0.36]; Tukey: P < 0.0001). Nevertheless, autonomous self-fertilized flowers significantly outperformed the ‘emasculated’ treated flowers (mean [LSE, USE]; fruit set: 0.4 [0.36, 0.44], fruit size: 20.5 cm^3^ [17.1, 24.5], proportion of fertilized achenes: 0.09 [0.07, 0.11]; Tukey: P < 0.0001; see Supplementary Section 2., Fig. 1). While ‘out-cross’ and ‘self-cross’ hand pollinations were equally successful at setting fruit (Tukey: P = 0.72), ‘self-cross’ pollinations produced significantly smaller fruit and with less proportion of fertilized achenes than ‘out-cross’ pollinations (Tukey: P < 0.0001; see Supplementary Section 2., Fig. 1. B and C). See Supplementary Section 2. Table 1. B for full model output.

#### Germination of achenes

Out of the 48 fruits we sampled achenes for germination, only ten produced achenes that germinated into cotyledon leaflets (50 out of a total of 127 fertilized achenes belonging to these ten fruits). All of these successful germinations happened with achenes that had been classified as ‘fertilized’ based on their shape and size. Achenes classified as ‘un-fertilized’ did not germinate. Germination rate was too low to test differences of germination rate across hand pollination treatments.

### Discussion

The hand pollination treatments confirm that *F. vesca* is self-compatible as there was no significant difference in fruit set and fruit size between self and outcross hand pollinations. Nevertheless, there is some evidence of inbreeding depression associated with selfing, as the proportion of fertilized achenes per fruit was higher in the outcross hand pollinations then in the self-cross hand pollinations. Therefore, while *F. vesca* is not heavily dependent on pollinators for reproductive success, it does gain a fitness advantage by the outcrossing achieved through pollination service. Furthermore, while *F. vesca* is fairly good at setting fruits through autonomous self-fertilization (80% fruit set), the proportion of fertilized achenes per autonomous fruit was significantly low (30%), highlighting the poor quality of these fruits. Emasculated flowers produced overall significantly lower fruit set, fruit size and proportion of fertilized achenes. Nevertheless, the fact that there was some fruit formation and achene fertilization opens up the possibility of either some wind pollination, apomixes, late emasculations or contamination by proximity to non-emasuclated flowers within an individual.

## Supplementary Section 2. Literature Cited

**Supplementary Section 2. Table 1.** Hand pollination treatment effects **(A)** Model description. **(B)** Model output.

| MODEL | RESPONSE VARIABLE | TRANSFORMATION ON RESPONSE VARIABLE | MODEL | DISTRIBUTION | FIXED EFFECTS* | RANDOM EFFECTS |
| --- | --- | --- | --- | --- | --- | --- |
| 1 | Fruit Set | None | Generalized Linear Mixed Model | Binomial | Hand Pollination Treatment<br>Plant Size** | Genotype<br>Block effect |
| 2 | Fruit Size | Log | Mixed Model | Normal | Hand Pollination Treatment<br>Plant Size**<br>Fruit Position | Genotype<br>Block effect |
| 3 | Proportion of Fertilized Achenes | Logit | Mixed Model | Normal | Hand Pollination Treatment<br>Plant Size**<br>Fruit Position | Genotype<br>Block effect |

\*Fixed effects did not include interactions; \*\*Plant size was standardized.

(B)
| Model | Response Variable | Fixed Effects | Df Num | Df Denom | F ratio | P value | Random Effects | Var ratio | Estimate | Std. Error | Wald p-value | % of Total |
| --- | --- | --- | --- | --- | --- | --- | --- | --- | --- | --- | --- | --- |
| 1 | Fruit Set | Hand Pollination Treatment | 3 | 1211 | 82.86 | < 0.0001 | Genotype |  | 0.26 | 0.14 | 0.055 |  |
|  |  | Plant Size | 1 | 414.7 | 2 | 0.16 | Block |  | 0.09 | 0.08 | 0.24 |  |
| 2 | Fruit Size | Hand Pollination Treatment | 3 | 823.4 | 86.66 | < 0.0001 | Genotype | 0.2 | 0.153 | 0.04 | < 0.0001 | 16.55 |
|  |  | Plant Size | 1 | 328.4 | 18.86 | < 0.0001 | Block | -0.0024 | -0.002 | 0.004 | 0.66 | 0 |
|  |  | Fruit Position* | 6 | 856.9 | 6.88 | < 0.0001 |  |  |  |  |  |  |
| 3 | Proportion of Fertilized Achenes | Hand Pollination Treatment | 3 | 739 | 201.63 | < 0.0001 | Genotype | 0.121 | 0.222 | 0.075 | 0.003 | 10.8 |
|  |  | Plant Size | 1 | 334.8 | 1.62 | 0.20 | Block | 0.0016 | 0.003 | 0.016 | 0.85 | 0.15 |
|  |  | Fruit Position* | 5 | 784.4 | 4.63 | 0.0004 |  |  |  |  |  |  |

\*Fruit position consisted originally of the three possible values within a *Fragaria vesca* inflorescence (primary, secondary or tertiary). However because values were averaged at the ramet level, fruit position values included in the model consisted of more than the three original values.

**Supplementary Section 2. Fig. 1.**
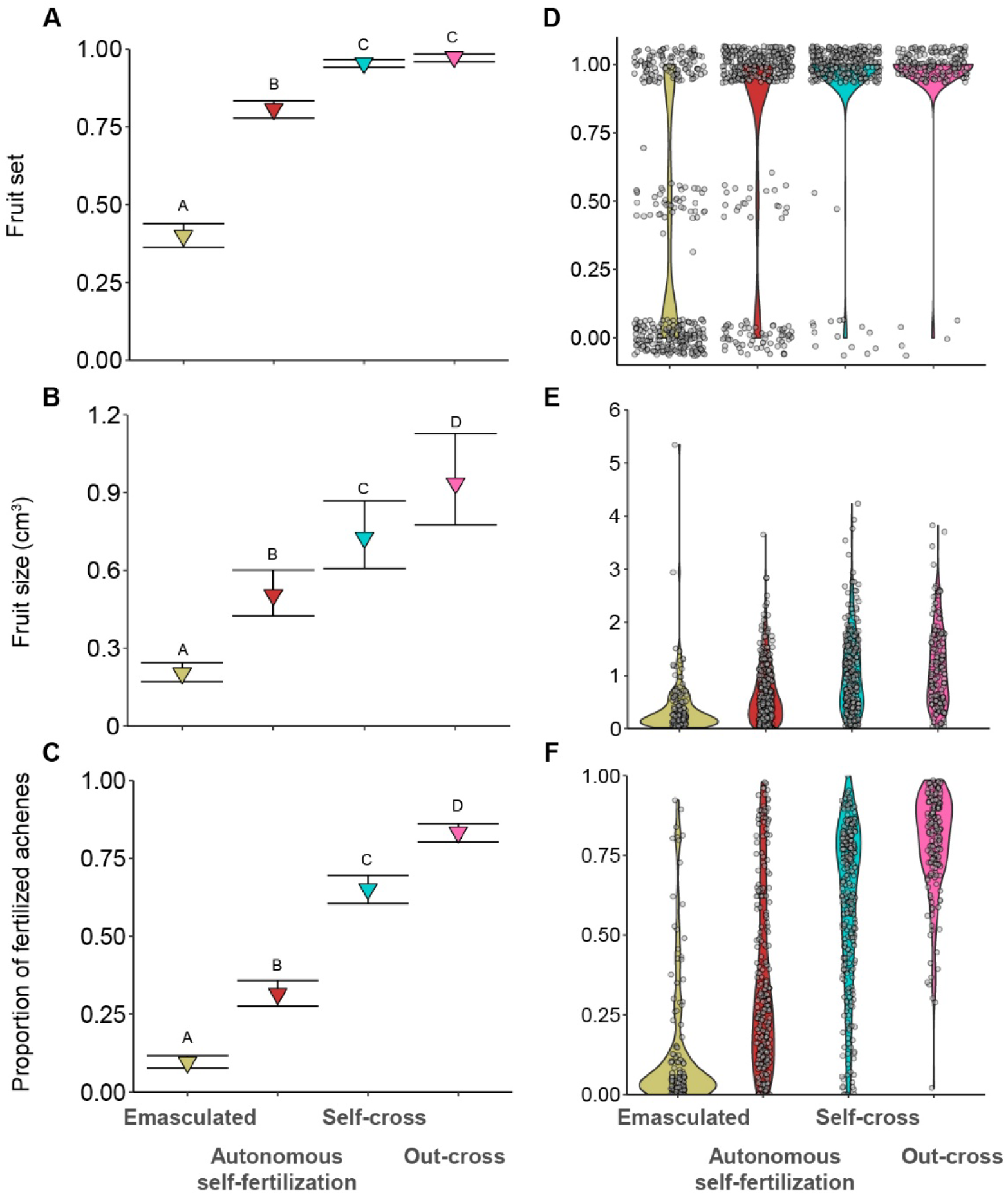
Flowers that manually received pollen either from the same genotype (Self-cross) or from a different genotype (Out-cross) produced higher quantity and quality of fruits. Hand-pollination treatment results across 121 *Fragaria vesca* genotypes, measured by (A, D) fruit set; (B, E) fruit size and (C, F) proportion of fertilized achenes. Panels A, B and C represent the marginal means estimated from the statistical models. Error bars are standard error. Letters denote significant differences for the Tukey HSD all pairwise comparison (P < 0.0001). Panels D, E and F are violin plots with the raw data points (grey circles). Each point on these violin plots represents a ramet. Panel D: Actual values ranged from zero to one. However, values go beyond this range due to the jitter effect used for plotting large number of data points. The presence of fruit set values other than zero is the result of sampling more than one fruit per ramet in many cases (see main text). Colors of symbols follow the corresponding hand pollination treatments labeled on the ‘x’ axis. All treatments were performed under a pollinator exclusion environment.

## Supplementary Section 3. Geographical coordinates of genotype samples

Supplementary Section 3. Geographical coordinates of genotype samples.
| Genotype | Region | Latitude | Longitude | Altitude (m.a.s.l) |
| --- | --- | --- | --- | --- |
| 10A | Sweden | 60.3738 | 17.5596667 | 106 |
| 12F | Sweden | 60.304 | 18.0151833 | 49 |
| 19A | Sweden | 60.15827 | 18.2141 | 21 |
| 19F | Sweden | 60.19762 | 18.1865667 | 218 |
| 1A | Sweden | 60.58625 | 17.4607833 | 19 |
| 23A | Sweden | 60.15597 | 17.4720834 | 167 |
| 35F | Sweden | 59.93182 | 16.8041 | 88 |
| 37A | Sweden | 59.92547 | 17.4046334 | 37 |
| 40A | Sweden | 59.75858 | 18.0445 | 22 |
| 43A | Sweden | 59.81657 | 17.34075 | 32 |
| 48F | Sweden | 59.63933 | 17.3085166 | 32 |
| 4A | Sweden | 60.49272 | 17.4682 | 27 |
| 6A | Sweden | 60.44652 | 17.7712667 | 17 |
| 8F | Sweden | 60.30762 | 18.5696667 | 25 |
| CRO1 | Croatia | 44.770848 | 15.652355 | 900 |
| CRO4 | Croatia | 45.838877 | 15.598735 | 320 |
| CRO5 | Croatia | 45.687132 | 15.381353 | 320 |
| CZ1 | Czech Republic | 50.86859 | 14.2971467 | 280 |
| ES1 | Spain | 43.3099 | -4.4697 | 70 |
| ES10 | Spain | 41.6176 | 0.62 | 185 |
| ES11 | Spain | 42.6712 | 0.9798 | 2190 |
| ES14 | Spain | 43.1425 | -4.938 | 974 |
| ES19 | Spain | 43.2504 | -5.9833 | 921 |
| ES21 | Spain | 37.2614 | -6.9447 | 34 |
| ES5 | Spain | 42.8117 | -1.6483 | 455 |
| ES7 | Spain | 42.7383 | -0.75 | 823 |
| ES8 | Spain | 42.6496 | 1.0347 | 2275 |
| ES9 | Spain | 42.7557 | -0.829 | 858 |
| FIN10 | Finland | 60.1949 | 21.3349 | 12 |
| FIN11 | Finland | 60.2 | 21.4 | 69 |
| FIN12 | Finland | 60.15 | 21.59 | 143 |
| FIN13 | Finland | 60.2292 | 25.0233 | 154 |
| FIN2 | Finland | 60.5367 | 21.8766 | 0 |
| FIN28 | Finland | 60.3431 | 25.6795 | 11 |
| FIN29 | Finland | 60.4775 | 24.8444 | 60 |
| FIN3 | Finland | 60.7262 | 21.9093 | 18 |
| FIN5 | Finland | 60.356 | 22.9651 | 28 |
| FIN50 | Finland | 60.1061 | 23.6782 | 32 |
| FIN51 | Finland | 60.0673 | 23.2879 | 21 |
| FIN53 | Finland | 60.2076 | 23.8066 | 34 |
| FIN55 | Finland | 60.4651 | 25.5254 | 205 |
| FIN6 | Finland | 59.8428 | 23.2446 | 12 |
| FIN7 | Finland | 60.3706 | 22.9796 | 38 |
| FR1 | France | 42.0396 | 9.0128926 | 1365 |
| FR4 | France | 48.42353 | 7.66326 | 157 |
| GER1 | Germany | 51.8236 | 11.2866 | 111 |
| GER2 | Germany | 51.1583 | 13.6814 | 181 |
| GER6 | Germany | 47.5857 | 10.5587 | 864 |
| GER7 | Germany | 50.58 | 13 | 576 |
| ICE11 | Iceland | 65.95007 | -19.487877 | 0 |
| ICE14 | Iceland | 63.84167 | -18.593056 | 283 |
| ICE17 | Iceland | 63.97232 | -16.839655 | 125 |
| ICE18 | Iceland | 65.83333 | -23.183333 | 0 |
| ICE19 | Iceland | 66.00507 | -16.49788 | 0 |
| ICE2 | Iceland | 63.99881 | -19.960389 | 0 |
| ICE5 | Iceland | 64.07347 | -19.843444 | 230 |
| ICE6 | Iceland | 64.52483 | -21.441028 | 174 |
| ICE9 | Iceland | 65.75618 | -19.538124 | 0 |
| IT16 | Italy | 45.7526 | 10.8644 | 1863 |
| IT19 | Italy | 46.54159 | 11.71958 | 1813 |
| IT21 | Italy | 46.0166 | 11.2653 | 585 |
| IT23 | Italy | 46.06118 | 11.2474 | 521 |
| IT24 | Italy | 46.22 | 11.26 | 1197 |
| IT6 | Italy | 46.2622 | 11.2709 | 1026 |
| IT7 | Italy | 46.275 | 11.2818 | 1268 |
| IT9 | Italy | 46.0137 | 11.0507 | 1529 |
| LIT1 | Lithuania | 54.7976 | 25.3484 | 131 |
| LIT2 | Lithuania | 54.8921 | 24.1482 | 46 |
| LIT5 | Lithuania | 55.0938 | 26.0724 | 189 |
| LIT6 | Lithuania | 54.986 | 25.8061 | 149 |
| ALTA1:6 | Norway | 69.9843 | 23.5210211 | 101 |
| ALTA2:4 | Norway | 69.9843 | 23.5210211 | 101 |
| NOR10 | Norway | 70.1848 | 23.3726 | 184 |
| NOR105 | Norway | 59.38179 | 10.690076 | 49 |
| NOR106 | Norway | 59.59135 | 5.1946 | 22 |
| NOR107 | Norway | 59.03252 | 10.170094 | 16 |
| NOR108 | Norway | 58.90031 | 9.647544 | 3 |
| NOR11 | Norway | 69.5248 | 18.3752 | 0 |
| NOR111 | Norway | 59.20078 | 10.5719 | 12 |
| NOR112 | Norway | 58.9323 | 9.53672 | 35 |
| NOR117 | Norway | 58.783 | 9.2872 | 85 |
| NOR118 | Norway | 68.7719 | 16.1878 | 24 |
| NOR119 | Norway | 59.0277 | 9.8772 | 74 |
| NOR120 | Norway | 58.49489 | 8.856994 | 0 |
| NOR121 | Norway | 59.79135 | 5.522733 | 14 |
| NOR122 | Norway | 59.77248 | 5.453716 | 56 |
| NOR123 | Norway | 59.04326 | 9.689636 | 55 |
| NOR127 | Norway | 63.34187 | 10.57585 | 161 |
| NOR13 | Norway | 69.5359 | 20.3805 | 30 |
| NOR131 | Norway | 59.20753 | 10.93096 | 24 |
| NOR133 | Norway | 59.16833 | 10.452577 | 12 |
| NOR134 | Norway | 59.24621 | 10.405009 | 57 |
| NOR137 | Norway | 63.33377 | 8.4582 | 9 |
| NOR14 | Norway | 69.5274 | 20.3724 | 0 |
| NOR141 | Norway | 59.14931 | 11.125367 | 30 |
| NOR144 | Norway | 64.14299 | 11.276353 | 80 |
| NOR147 | Norway | 62.91398 | 8.184851 | 45 |
| NOR150 | Norway | 58.91914 | 5.99499 | 0 |
| NOR153 | Norway | 62.28653 | 6.95902 | 168 |
| NOR16 | Norway | 69.4451 | 20.949 | 104 |
| NOR162 | Norway | 59.38285 | 10.652894 | 11 |
| NOR164 | Norway | 63.69261 | 10.287903 | 0 |
| NOR17 | Norway | 69.4256 | 20.9715 | 144 |
| NOR172 | Norway | 58.6469 | 9.1179 | 43 |
| NOR174 | Norway | 63.13235 | 10.326228 | 136 |
| NOR177 | Norway | 68.80165 | 17.2212 | 77 |
| NOR2 | Norway | 69.9395 | 23.0964 | 29 |
| NOR20 | Norway | 70.0227 | 22.0173 | 0 |
| NOR3 | Norway | 69.9396 | 23.0971 | 19 |
| NOR30 | Norway | 69.78389 | 18.538889 | 77 |
| NOR8 | Norway | 70.0324 | 23.4012 | 83 |
| NOR9 | Norway | 70.1585 | 23.2781 | 9 |
| OLAND1:1 | Sweden | 57.35297 | 17.1258928 | 10 |
| POR2 | Portugal | 38.7984 | -9.38811 | 198 |
| UK1 | Great Britain | 57.8954 | -5.1613 | 11 |
| UK11 | Great Britain | 54.7331 | -3.2105 | 164 |
| UK12 | Great Britain | 50.2451 | -3.77663 | 31 |
| UK2 | Great Britain | 54.6014 | -3.3197 | 86 |
| UK4 | Great Britain | 51.8757 | -4.9392 | 85 |
| UK6 | Great Britain | 57.5667 | -3.8833 | 24 |
| UK9 | Great Britain | 51.4845 | -1.0261 | 82 |

\*All sampling locations were at least a few kilometers apart from each other, except for a two samples from northern Norway (NOR2 and NOR3) with a sampling distance of about 20 meters.

## Supplementary Section 4. Subset of genotypes included in inbreeding depression analysis

**Supplementary Section 4. Table 1.** Subset of genotypes (34 out of 121) included in inbreeding depression analyses. Table includes genotype abbreviation, number of replicates per genotype (i.e. number of ramets), and geographical coordinates.

| **Genotype** | **Replicates per Genotype** | **Latitude** | **Longitude** |
| --- | --- | --- | --- |
| **10A** | 3 | 60.3738 | 17.55967 |
| **12F** | 2 | 60.304 | 18.01518 |
| **19F** | 2 | 60.19762 | 18.18657 |
| **1A** | 3 | 60.58625 | 17.46078 |
| **35F** | 2 | 59.93182 | 16.8041 |
| **43A** | 2 | 59.81657 | 17.34075 |
| **4A** | 2 | 60.49272 | 17.4682 |
| **CRO1** | 2 | 44.7708 | 15.65235 |
| **CRO4** | 2 | 45.8388 | 15.598735 |
| **FIN3** | 2 | 60.7262 | 21.9093 |
| **FIN5** | 3 | 60.356 | 22.9651 |
| **FR1** | 2 | 42.0396 | 9.012893 |
| **ICE14** | 3 | 63.84167 | -18.5931 |
| **ICE5** | 3 | 64.07347 | -19.8434 |
| **ICE6** | 2 | 64.52483 | -21.441 |
| **IT16** | 3 | 45.7526 | 10.8644 |
| **IT21** | 2 | 46.0166 | 11.2653 |
| **IT23** | 3 | 46.06118 | 11.2474 |
| **IT6** | 4 | 46.2622 | 11.2709 |
| **IT9** | 3 | 46.0137 | 11.0507 |
| **LIT1** | 2 | 54.7976 | 25.3484 |
| **LIT2** | 2 | 54.8921 | 24.1482 |
| **LIT5** | 3 | 55.0938 | 26.0724 |
| **LIT6** | 3 | 54.986 | 25.8061 |
| **NOR10** | 2 | 70.1848 | 23.3726 |
| **NOR105** | 2 | 59.38179 | 10.69008 |
| **NOR106** | 2 | 59.59135 | 5.1946 |
| **NOR107** | 4 | 59.03252 | 10.17009 |
| **NOR11** | 3 | 69.5248 | 18.3752 |
| **NOR111** | 2 | 59.20078 | 10.5719 |
| **NOR112** | 2 | 58.9323 | 9.53672 |
| **NOR144** | 3 | 64.14299 | 11.27635 |
| **NOR9** | 2 | 70.1585 | 23.2781 |
| **UK12** | 2 | 50.2451 | -3.77663 |

**Supplementary Section 4. Fig. 1.**
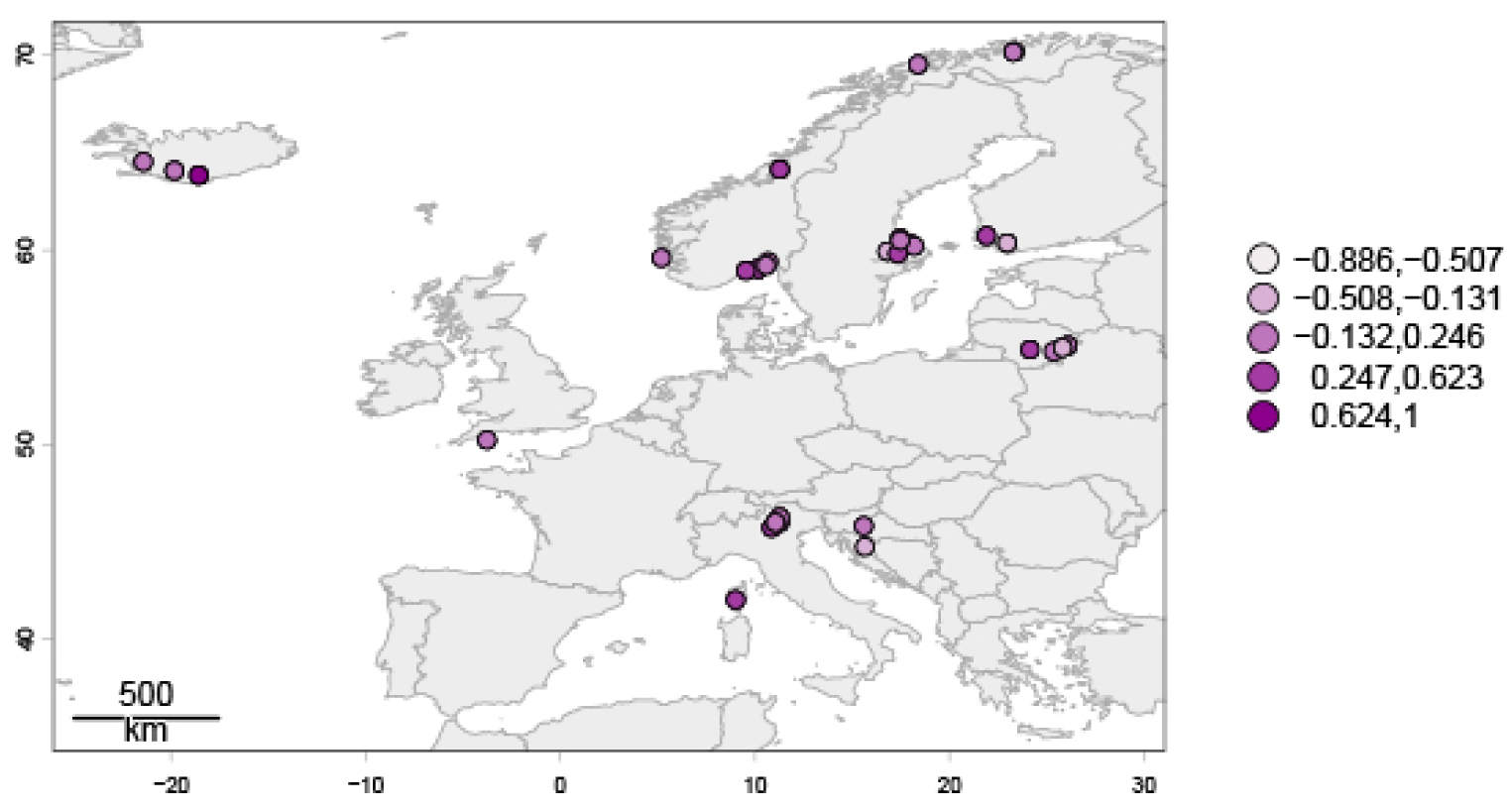
Geographical distribution of genotypes included for the relative performance (RP) analysis. Colour coded based on RP values. Darker colours indicate more positive values of RP (i.e. more inbreeding depression).

## Supplementary Section 5. Summary Statistics

**Supplementary Section 5. Table 1.**
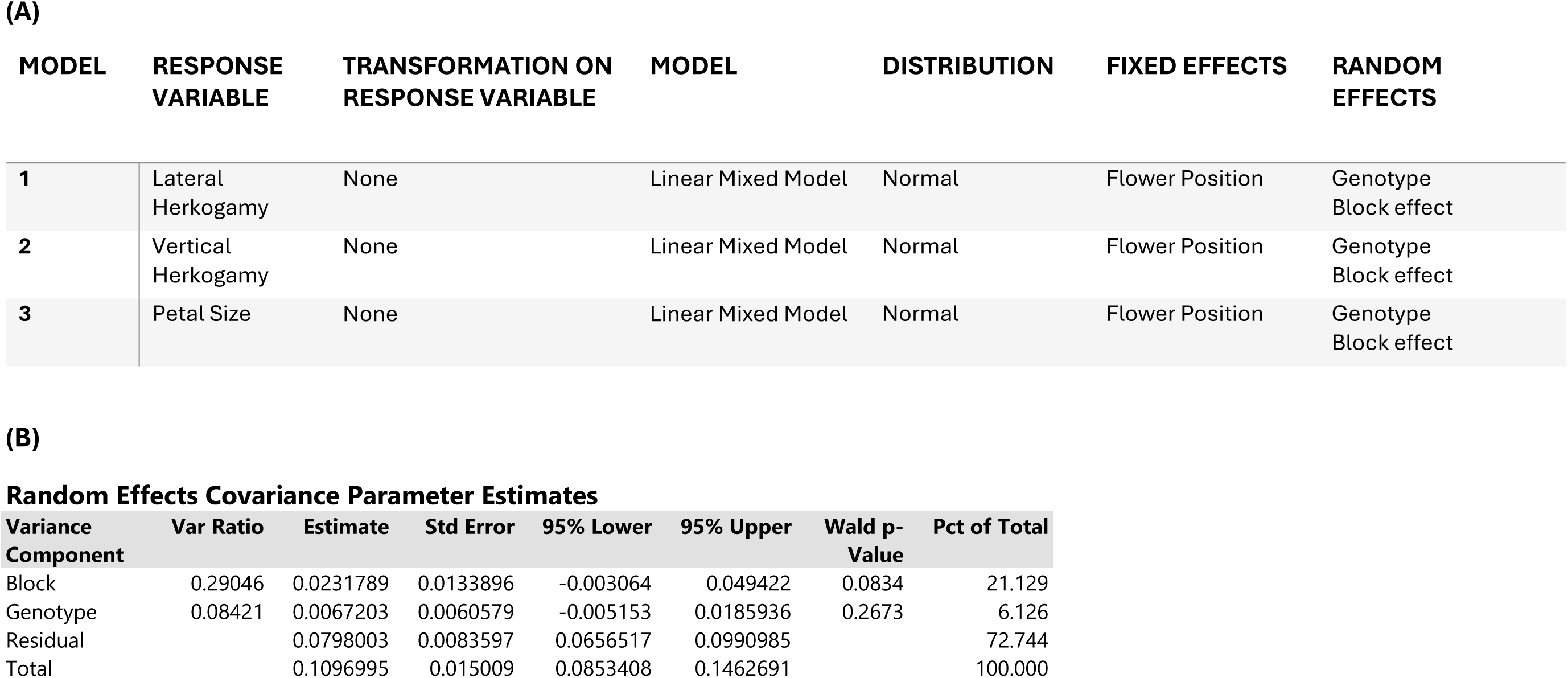
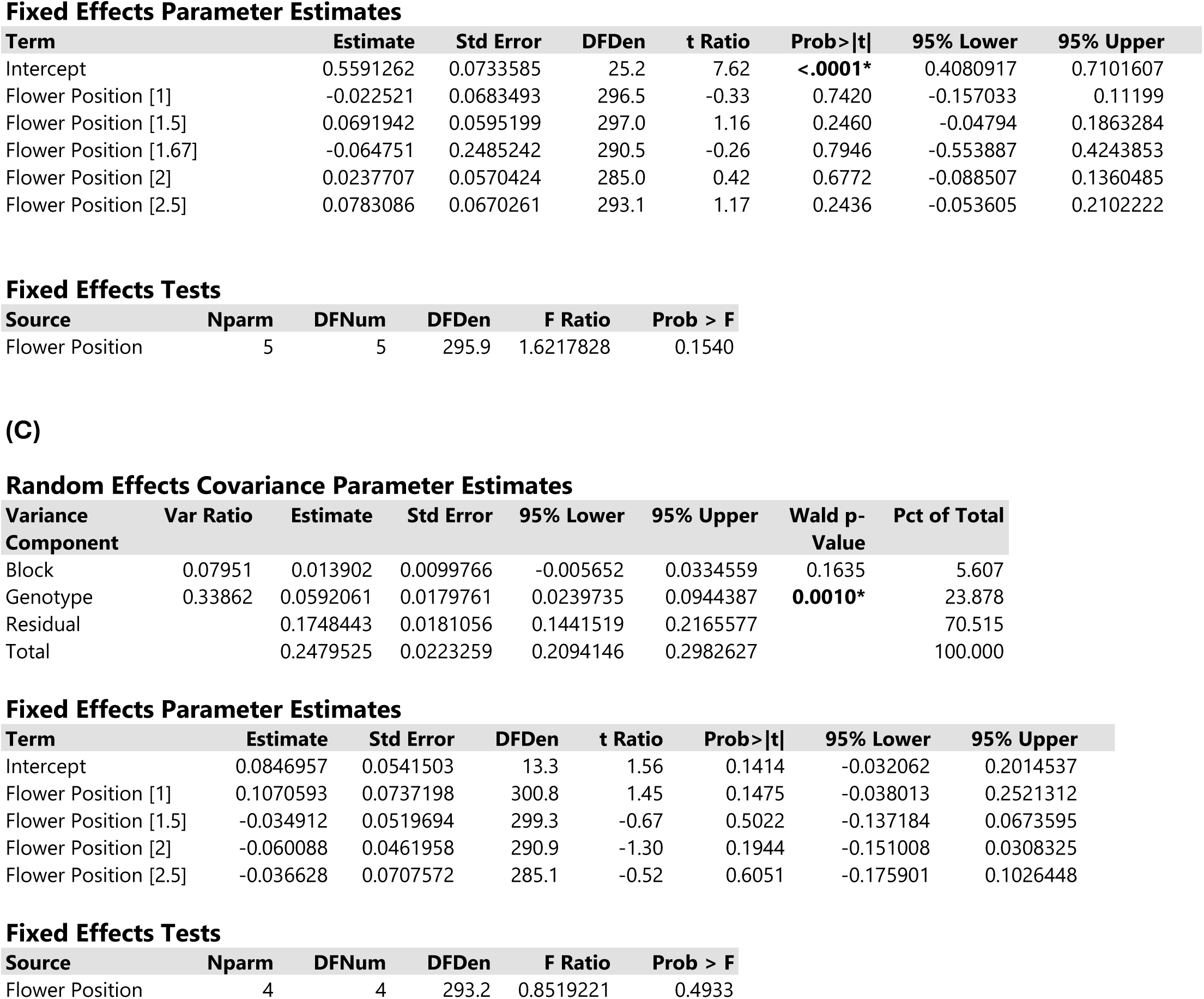
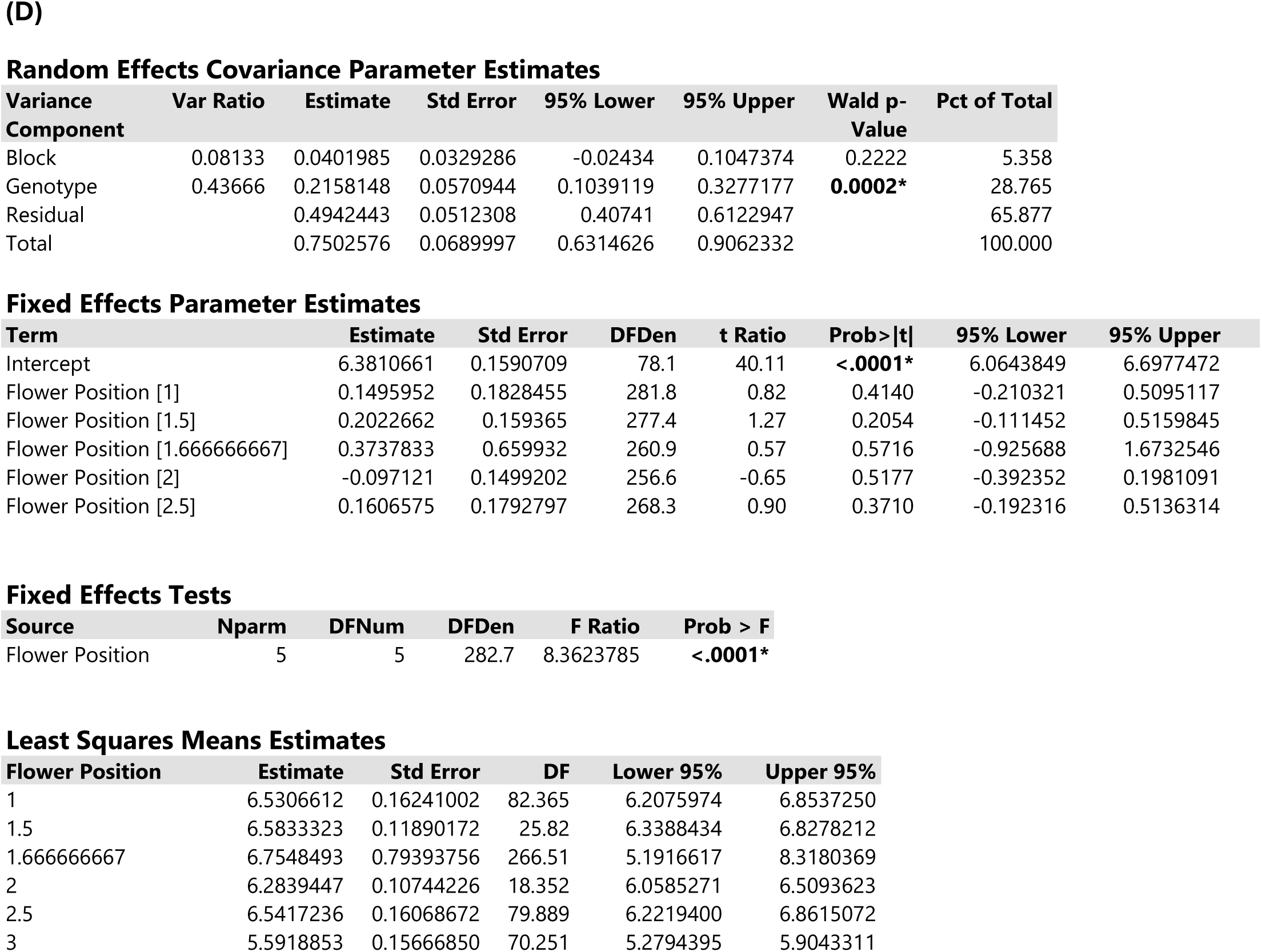
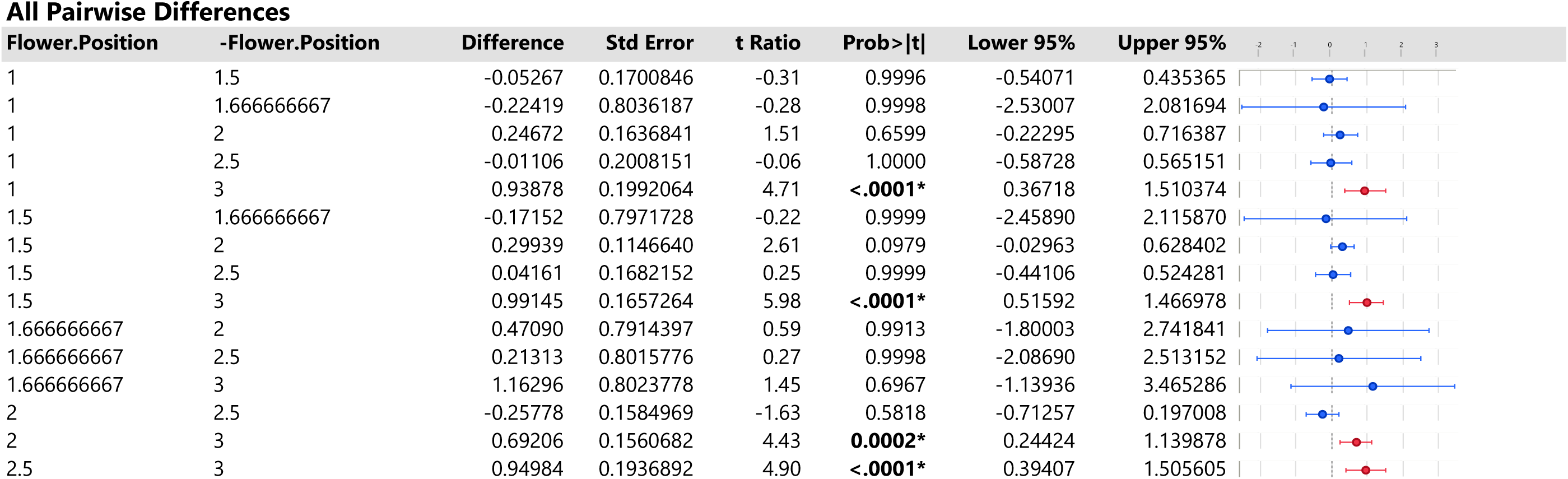
Flower position within the inflorescence does not affect values of lateral and vertical herkogamy. Tertiary flowers, however, have smaller petals than primary and secondary flowers. Descriptions of all three models **(A)** and model output for lateral herkogamy **(B)**, vertical herkogamy **(C)** and petal size **(D)**. Flower position consisted originally of the three possible values within a *Fragaria vesca* inflorescence (primary, secondary or tertiary). However, because values were averaged at the ramet level, flower position values included in the model consisted of more than the three original values. Values in bold are statistically significant.

**Supplementary Section 5. Table 2.** Fruit position within the inflorescence does not explain variation across traits measured on autonomous fruits. **(A)**. Model description. Model output for **(B)** fruit size and **(C) the** proportion of fertilized achenes. Fruit position consisted originally of the three possible values within a *Fragaria vesca* inflorescence (primary, secondary or tertiary). However, since values were averaged at the ramet level, fruit position values included in the model consisted of more than the three original values. Values in bold are statistically significant.

| MODEL | RESPONSE VARIABLE | TRANSFORMATION ON RESPONSE VARIABLE | MODEL | DISTRIBUTION | FIXED EFFECTS | RANDOM EFFECTS |
| --- | --- | --- | --- | --- | --- | --- |
| 1 | Fruit Size | Log | Linear Mixed Model | Normal | Fruit Position | Genotype<br>Block effect |
| 2 | Proportion of Fertilized Achenes | Logit | Linear Mixed Model | Normal | Fruit Position | Genotype<br>Block effect |

**(B)** **Random Effects Covariance Parameter Estimates**
| Variance Component | Var Ratio | Estimate | Std Error | 95% Lower | 95% Upper | Wald p-Value | Pct of Total |
| --- | --- | --- | --- | --- | --- | --- | --- |
| Block | 0.00847 | 0.0064781 | 0.0182851 | -0.02936 | 0.0423163 | 0.7231 | 0.651 |
| Genotype | 0.29168 | 0.2231886 | 0.0886268 | 0.0494832 | 0.3968939 | **0.0118\*** | **22.434** |
| Residual |  | 0.765188 | 0.0867353 | 0.6200505 | 0.9682947 |  | 76.915 |
| Total |  | 0.9948547 | 0.0881516 | 0.8423975 | 1.1930371 |  | 100.000 |

**Fixed Effects Parameter Estimates**
| Term | Estimate | Std Error | DFDen | t Ratio | Prob>\|t\| | 95% Lower | 95% Upper |
| --- | --- | --- | --- | --- | --- | --- | --- |
| Intercept | 6.1962665 | 0.1384072 | 100.0 | 44.77 | <.0001* | 5.9216695 | 6.4708635 |
| Fruit.Position[1] | 0.2542133 | 0.1909686 | 269.5 | 1.33 | 0.1843 | -0.121767 | 0.6301931 |
| Fruit.Position[1.5] | -0.15357 | 0.2010836 | 257.5 | -0.76 | 0.4457 | -0.549548 | 0.2424082 |
| Fruit.Position[2] | 0.0358429 | 0.14678 | 277.0 | 0.24 | 0.8073 | -0.253103 | 0.3247888 |
| Fruit.Position[2.333333333] | 0.2279016 | 0.5763512 | 270.3 | 0.40 | 0.6928 | -0.906806 | 1.3626092 |
| Fruit.Position[2.5] | -0.10828 | 0.1996419 | 242.3 | -0.54 | 0.5881 | -0.501535 | 0.2849752 |

**Fixed Effects Tests**
| Source | Nparm | DFNum | DFDen | F Ratio | Prob > F |
| --- | --- | --- | --- | --- | --- |
| Fruit Position | 5 | 5 | 252.3 | 1.6099987 | 0.1578 |

**(C)**55 **Random Effects Covariance Parameter Estimates**
| Variance Component | Var Ratio | Estimate | Std Error | 95% Lower | 95% Upper | Wald p-Value | Pct of Total |
| --- | --- | --- | --- | --- | --- | --- | --- |
| Block | 0.03216 | 0.0834836 | 0.1155054 | -0.142903 | 0.30987 | 0.4698 | 3.082 |
| Genotype | 0.01143 | 0.0296807 | 0.2007266 | -0.363736 | 0.4230976 | 0.8824 | 1.096 |
| Residual |  | 2.5959469 | 0.299792 | 2.0957004 | 3.3004719 |  | 95.823 |
| Total |  | 2.7091112 | 0.2479938 | 2.2819011 | 3.2693173 |  | 100.000 |

**Fixed Effects Parameter Estimates**
| Term | Estimate | Std Error | DFDen | t Ratio | Prob>\|t\| | 95% Lower | 95% Upper |
| --- | --- | --- | --- | --- | --- | --- | --- |
| Intercept | -0.63444 | 0.2462079 | 36.9 | -2.58 | **0.0141\*** | -1.13335 | -0.13553 |
| Fruit.Position[1] | -0.814396 | 0.3346266 | 252.1 | -2.43 | **0.0156\*** | -1.473416 | -0.155377 |
| Fruit.Position[1.5] | -0.48075 | 0.3752855 | 249.2 | -1.28 | 0.2014 | -1.219887 | 0.2583861 |
| Fruit.Position[2] | -0.108447 | 0.2552646 | 250.6 | -0.42 | 0.6713 | -0.611184 | 0.3942902 |
| Fruit.Position[2.333333333] | 1.5458957 | 0.9799525 | 250.1 | 1.58 | 0.1159 | -0.384115 | 3.4759067 |
| Fruit.Position[2.5] | -0.171858 | 0.3747613 | 244.4 | -0.46 | 0.6469 | -0.910031 | 0.5663158 |

**Fixed Effects Tests**
| Source | Nparm | DFNum | DFDen | F Ratio | Prob > F |
| --- | --- | --- | --- | --- | --- |
| Fruit Position | 5 | 5 | 248.6 | 1.7123615 | 0.1323 |

**Supplementary Section 5. Table. 3.**
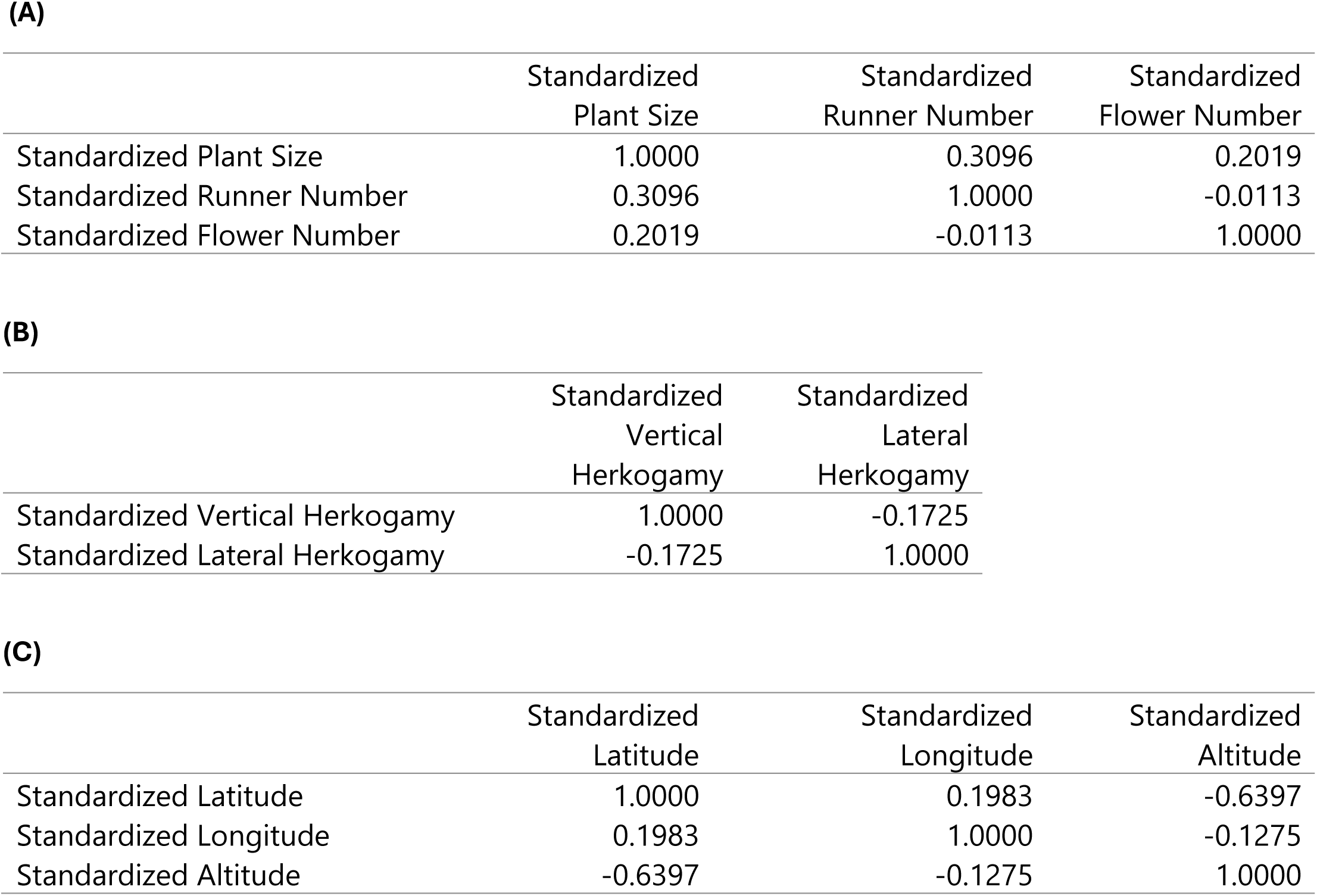
Pearson correlation plot for predictors included in models testing the **(A)** allocation between selfing and cloning, **(B)** relationship between floral traits and the capacity to autonomously self-fertilize and **(C)** geographical coordinates and elevation. All correlations are below 0.7 and hence no indication of multicolinearity between predictor variables included in these models (Dormann et al., 2013).

**Supplementary Section 5. Table 4.**
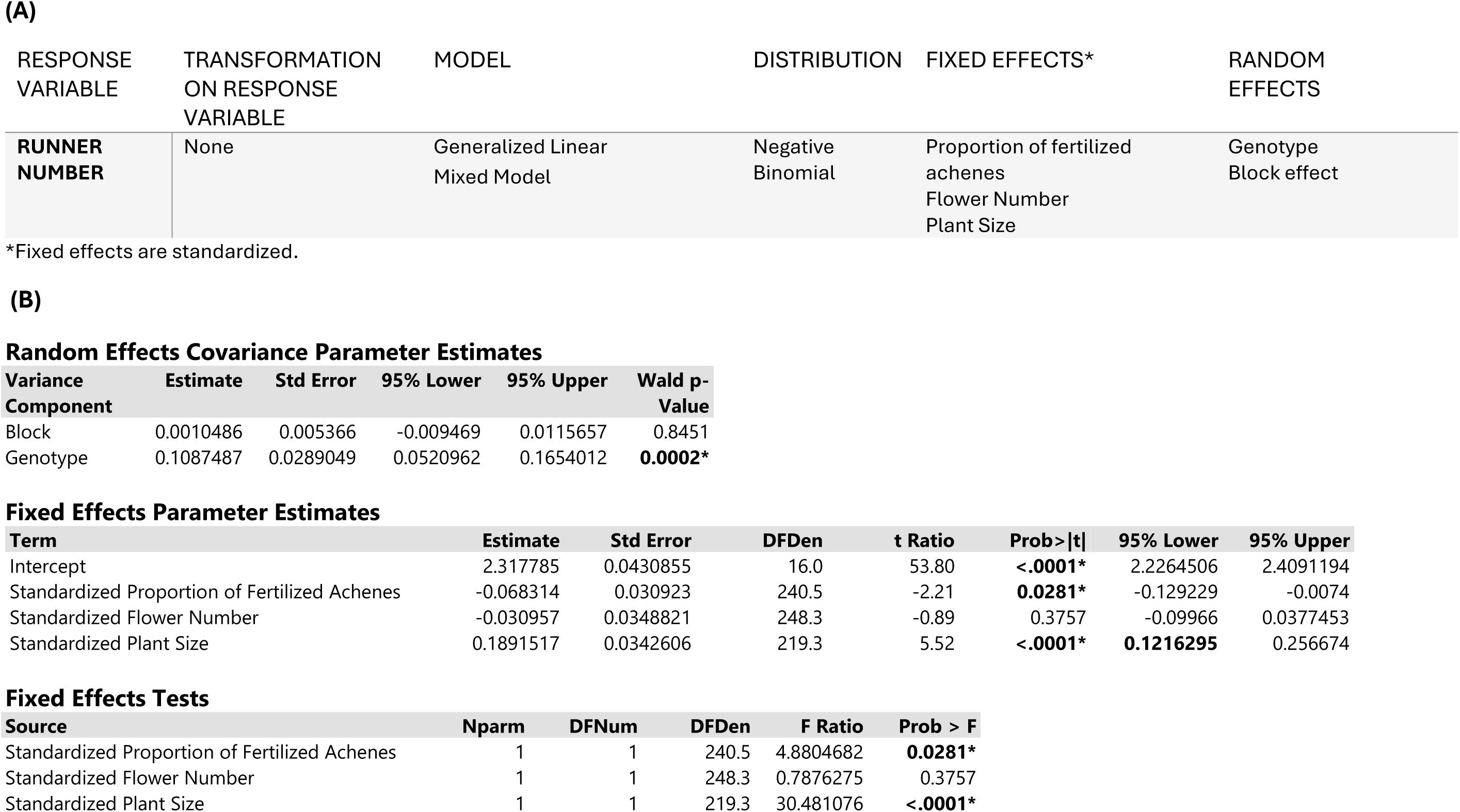
Analysis details describing the reproductive assurance allocation / trade off. **(A)** Model description. **(B)** Model output. Values in bold are statistically significant.

**Supplementary Section 5. Table 5.** Analyses details for reproductive assurance allocation / trade off – Inverted model. **(A)** Model description. **(B)** Model output. Values in bold are statistically significant.

| RESPONSE<br>VARIABLE | TRANSFORMATION<br>ON RESPONSE<br>VARIABLE | MODEL | DISTRIBUTION | FIXED EFFECTS* | RANDOM<br>EFFECTS |
| --- | --- | --- | --- | --- | --- |
| **PROPORTION**<br>**OF FERTILIZED**<br>**ACHENES** | Logit | Mixed Model | Normal | Runner Number<br>Flower Number<br>Plant Size | Genotype<br>Block effect |

| Variance<br>Component | Var Ratio | Estimate | Std Error | 95% Lower | 95% Upper | Wald p-Value | Pct of Total |
| --- | --- | --- | --- | --- | --- | --- | --- |
| Block | 0.01672 | 0.044976 | 0.0984239 | -0.147931 | 0.2378833 | 0.6477 | 1.645 |
| Genotype | -0.0189 | -0.050847 | 0.1975419 | -0.438022 | 0.3363281 | 0.7969 | 0.000 |
| Residual |  | 2.6896596 | 0.3124639 | 2.1687394 | 3.4248099 |  | 98.355 |
| Total |  | 2.7346356 | 0.3206195 | 2.2008709 | 3.4903296 |  | 100.000 |

**Fixed Effects Parameter Estimates**
| Term | Estimate | Std Error | DFDen | t Ratio | Prob>\|t\| | 95% Lower | 95% Upper |
| --- | --- | --- | --- | --- | --- | --- | --- |
| Intercept | -0.824132 | 0.1250976 | 3.7 | -6.59 | **0.0036** | -1.183334 | -0.46493 |
| Standardized Plant Size | 0.2467552 | 0.1163127 | 204.5 | 2.12 | **0.0351** | 0.0174292 | 0.4760812 |
| Standardized Flower Number | -0.127473 | 0.1208019 | 216.6 | -1.06 | 0.2925 | -0.365571 | 0.1106245 |
| Standardized Runner Number | -0.249613 | 0.1088963 | 213.4 | -2.29 | **0.0229** | -0.464263 | -0.034963 |

**Fixed Effects Tests**
| Source | Nparm | DFNum | DFDen | F Ratio | Prob > F |
| --- | --- | --- | --- | --- | --- |
| Standardized Plant Size | 1 | 1 | 204.5 | 4.500683 | **0.0351** |
| Standardized Flower Number | 1 | 1 | 216.6 | 1.1134988 | 0.2925 |
| Standardized Runner Number | 1 | 1 | 213.4 | 5.2542225 | **0.0229** |

**Supplementary Section 5. Table 6.** Analyses details for effects of lateral and vertical herkogamy on capacity to self-fertilize. **(A)** Model description. **(B)** Model output. Values in bold are statistically significant.

| RESPONSE<br>VARIABLE | TRANSFORMATION<br>ON RESPONSE<br>VARIABLE | MODEL | DISTRIBUTION | FIXED EFFECTS* | RANDOM<br>EFFECTS |
| --- | --- | --- | --- | --- | --- |
| **PROPORTION**<br>**OF FERTILIZED**<br>**ACHENES** | Logit | Linear Mixed Model | Normal | Lateral Herkogamy<br>Vertical Herkogamy | Genotype<br>Block effect |

| Variance<br>Component | Var Ratio | Estimate | Std Error | 95% Lower | 95% Upper | Wald p-Value | Pct of Total |
| --- | --- | --- | --- | --- | --- | --- | --- |
| Block | 0.01028 | 0.0279507 | 0.0927024 | -0.153743 | 0.209644 | 0.7630 | 1.017 |
| Genotype | -0.0482 | -0.131039 | 0.2285297 | -0.578949 | 0.3168714 | 0.5664 | 0.000 |
| Residual |  | 2.7199512 | 0.3529246 | 2.1419203 | 3.5695454 |  | 98.983 |
| Total |  | 2.7479019 | 0.3580551 | 2.1618841 | 3.6106495 |  | 100.000 |

**Fixed Effects Parameter Estimates**
| Term | Estimate | Std Error | DFDen | t Ratio | Prob>\|t\| | 95% Lower | 95% Upper |
| --- | --- | --- | --- | --- | --- | --- | --- |
| Intercept | -0.881204 | 0.1220627 | 3.6 | -7.22 | **0.0029\*** | -1.235958 | -0.52645 |
| Standardized Vertical Herkogamy | 0.0049451 | 0.1202969 | 175.1 | 0.04 | 0.9673 | -0.232474 | 0.2423639 |
| Standardized Lateral Herkogamy | -0.299117 | 0.115632 | 118.2 | -2.59 | **0.0109\*** | -0.528095 | -0.070139 |
| (Stand. [Vertical Herkogamy]+0.02553)* (Stand. [Lateral Herkogamy] +0.00936) | -0.00971 | 0.1196501 | 202.7 | -0.08 | 0.9354 | -0.245628 | 0.2262078 |

**Fixed Effects Tests**
| Source | Nparm | DFNum | DFDen | F Ratio | Prob > F |
| --- | --- | --- | --- | --- | --- |
| Standardized Vertical Herkogamy | 1 | 1 | 175.1 | 0.0016898 | 0.9673 |
| Standardized Lateral Herkogamy | 1 | 1 | 118.2 | 6.6915436 | **0.0109\*** |
| Standardized Vertical Herkogamy*Standardized Lateral Herkogamy | 1 | 1 | 202.7 | 0.0065863 | 0.9354 |

**Supplementary Section 5. Table 7.** The effect of inbreeding depression (measured as relative performance) on the capacity to autonomously self-fertilize. **(A)** Model description. **(B)** Model output. Values in bold are statistically significant.

| RESPONSE<br>VARIABLE | TRANSFORMATION<br>ON RESPONSE<br>VARIABLE | MODEL | DISTRIBUTION | FIXED EFFECTS | RANDOM<br>EFFECTS |
| --- | --- | --- | --- | --- | --- |
| **PROPORTION**<br>**OF FERTILIZED**<br>**ACHENES FOR**<br>**AUTONOMOUS**<br>**FRUITS** | Logit | Linear Mixed Model | Normal | Relative Performance | Genotype<br>Block effect |

| Variance<br>Component | Var Ratio | Estimate | Std Error | 95% Lower | 95% Upper | Wald p-Value | Pct of Total |
| --- | --- | --- | --- | --- | --- | --- | --- |
| Genotype | 0.05638 | 0.1239045 | 0.3915177 | -0.643456 | 0.891265 | 0.7516 | 5.337 |
| Block | -0.0316 | -0.06938 | 0.063171 | -0.193193 | 0.0544332 | 0.2721 | 0.000 |
| Residual |  | 2.1976434 | 0.5090263 | 1.4626819 | 3.6703624 |  | 94.663 |
| Total |  | 2.3215478 | 0.3741354 | 1.732886 | 3.2725256 |  | 100.000 |

**Fixed Effects Parameter Estimates**
| Term | Estimate | Std Error | DFDen | t Ratio | Prob>\|t\| | 95% Lower | 95% Upper |
| --- | --- | --- | --- | --- | --- | --- | --- |
| Intercept | -0.883787 | 0.1823456 | 8.5 | -4.85 | **0.0011\*** | -1.300237 | -0.467337 |
| Relative<br>Performance | -0.03605 | 0.6083172 | 69.3 | -0.06 | 0.9529 | -1.249524 | 1.1774238 |

**Fixed Effects Tests**
| Source | Nparm | DFNum | DFDen | F Ratio | Prob > F |
| --- | --- | --- | --- | --- | --- |
| Relative<br>Performance | 1 | 1 | 69.3 | 0.003512 | 0.9529 |

**Supplementary Section 5. Table 8.** Capacity to self- fertilize across geographical coordinates and altitude. **(A)** Model description. **(B)** Model output. Values in bold are statistically significant.

| RESPONSE<br>VARIABLE | TRANSFORMATION<br>ON RESPONSE<br>VARIABLE | MODEL | DISTRIBUTION | FIXED EFFECTS* | RANDOM EFFECTS |
| --- | --- | --- | --- | --- | --- |
| **PROPORTION**<br>**OF**<br>**FERTILIZED**<br>**ACHENES** | Logit | Linear Mixed Model | Normal | Latitude<br>Longitude<br>Altitude | Genotype<br>Block |

| Variance<br>Component | Var Ratio | Estimate | Std Error | 95% Lower | 95% Upper | Wald p-Value | Pct of Total |
| --- | --- | --- | --- | --- | --- | --- | --- |
| Block | 0.02938 | 0.0754365 | 0.1083939 | -0.137012 | 0.2878847 | 0.4865 | 2.773 |
| Genotype | 0.03021 | 0.0775665 | 0.1981888 | -0.310876 | 0.4660093 | 0.6955 | 2.851 |
| Residual |  | 2.5672371 | 0.290272 | 2.0813323 | 3.2466403 |  | 94.375 |
| Total |  | 2.7202401 | 0.2470993 | 2.2941676 | 3.2777865 |  | 100.000 |

**Fixed Effects Parameter Estimates**
| Term | Estimate | Std Error | DFDen | t Ratio | Prob>\|t\| | 95% Lower | 95% Upper |
| --- | --- | --- | --- | --- | --- | --- | --- |
| Intercept | -0.81278 | 0.140949 | 5.2 | -5.77 | **0.0020\*** | -1.171497 | -0.454063 |
| Standardized Latitude | -0.320242 | 0.1429251 | 96.4 | -2.24 | **0.0273\*** | -0.603933 | -0.036551 |
| Standardized Longitude | 0.0368475 | 0.1043456 | 81.7 | 0.35 | 0.7249 | -0.170739 | 0.2444345 |
| Standardized Altitude | -0.317212 | 0.1449286 | 88.3 | -2.19 | **0.0313\*** | -0.605214 | -0.02921 |

**Fixed Effects Tests**
| Source | Nparm | DFNum | DFDen | F Ratio | Prob > F |
| --- | --- | --- | --- | --- | --- |
| Standardized Latitude | 1 | 1 | 96.4 | 5.020417 | **0.0273\*** |
| Standardized Longitude | 1 | 1 | 81.7 | 0.1247006 | 0.7249 |
| Standardized Altitude | 1 | 1 | 88.3 | 4.7906148 | **0.0313\*** |

**Supplementary Section 5. Table 9.** Propensity to clone across geographical coordinates and altitude. **(A)** Model description. **(B)** Model output. Values in bold are statistically significant.

| RESPONSE<br>VARIABLE | TRANSFORMATION<br>ON RESPONSE<br>VARIABLE | MODEL | DISTRIBUTION | FIXED EFFECTS* | RANDOM EFFECTS |
| --- | --- | --- | --- | --- | --- |
| **RUNNER**<br>**NUMBER** | None | Generalized Linear Mixed<br>Model | Negative<br>Binomial | Latitude<br>Longitude<br>Altitude | Genotype<br>Block |

| Variance<br>Component | Estimate | Std Error | 95% Lower | 95% Upper | Wald p-Value |
| --- | --- | --- | --- | --- | --- |
| Block | 0.0059476 | 0.0067342 | -0.007251 | 0.0191464 | 0.3771 |
| Genotype | 0.1141248 | 0.0263108 | 0.0625566 | 0.1656931 | **<.0001\*** |

**Fixed Effects Parameter Estimates**
| Term | Estimate | Std Error | DFDen | t Ratio | Prob>\|t\| | 95% Lower | 95% Upper |
| --- | --- | --- | --- | --- | --- | --- | --- |
| Intercept | 2.310065 | 0.0473429 | 15.7 | 48.79 | **<.0001\*** | 2.2095688 | 2.4105612 |
| Standardized Latitude | -0.044169 | 0.0510443 | 105.9 | -0.87 | 0.3888 | -0.145371 | 0.0570328 |
| Standardized Longitude | 0.0906936 | 0.0411994 | 108.1 | 2.20 | **0.0298\*** | 0.0090301 | 0.172357 |
| Standardized Altitude | -0.057737 | 0.0505076 | 108.0 | -1.14 | 0.2555 | -0.157852 | 0.0423774 |

**Fixed Effects Tests**
| Source | Nparm | DFNum | DFDen | F Ratio | Prob > F |
| --- | --- | --- | --- | --- | --- |
| Standardized Latitude | 1 | 1 | 105.9 | 0.7487632 | 0.3888 |
| Standardized Longitude | 1 | 1 | 108.1 | 4.8458653 | **0.0298\*** |
| Standardized Altitude | 1 | 1 | 108.0 | 1.3067809 | 0.2555 |

**Supplementary Section 5. Table 10.** The relationship between geographical coordinates and reproductive traits involved in pollinator attraction. **(A)** Model description. **(B)** Model output for petal size. **(C)** model output for inflorescence exsertion and **(D)** model output for flower number. Values in bold are statistically significant.

| MODEL | RESPONSE VARIABLE | TRANSFORMATION ON RESPONSE VARIABLE | MODEL | DISTRIBUTION | FIXED EFFECTS* | RANDOM EFFECT |
| --- | --- | --- | --- | --- | --- | --- |
| **1** | Petal Size | None** | Linear Mixed Model | Normal | Latitude<br>Longitude<br>Altitude<br>Latitude*Longitude | Genotype<br>Block |
| **2** | Inflorescence Exsertion | None | Linear Mixed Model | Normal | Latitude<br>Longitude<br>Altitude | Genotype<br>Block |
| **3** | Flower Number | None | Generalized Linear Mixed Model | Negative Binomial | Latitude<br>Longitude<br>Altitude | Genotype<br>Block |

| Variance Component | Var Ratio | Estimate | Std Error | 95% Lower | 95% Upper | Wald p-Value | Pct of Total |
| --- | --- | --- | --- | --- | --- | --- | --- |
| Block | 0.48756 | 0.0479727 | 0.0266879 | -0.004335 | 0.1002801 | 0.0722 | 17.994 |
| Genotype | 1.22195 | 0.1202318 | 0.0215445 | 0.0780054 | 0.1624582 | **<.0001\*** | 45.099 |
| Residual |  | 0.0983933 | 0.0102119 | 0.0810875 | 0.1219296 |  | 36.907 |
| Total |  | 0.2665978 | 0.0347618 | 0.2097108 | 0.3503705 |  | 100.000 |

**Fixed Effects Parameter Estimates**
| Term | Estimate | Std Error | DFDen | t Ratio | Prob>\|t\| | 95% Lower | 95% Upper |
| --- | --- | --- | --- | --- | --- | --- | --- |
| Intercept | -0.022634 | 0.0827291 | 10.5 | -0.27 | 0.7897 | -0.205721 | 0.1604525 |
| Standardized Latitude | 0.1064852 | 0.0499843 | 118.5 | 2.13 | **0.0352\*** | 0.007507 | 0.2054635 |
| Standardized Longitude | -0.072228 | 0.0411071 | 114.5 | -1.76 | 0.0816 | -0.153658 | 0.0092009 |
| Standardized Altitude | 0.0772277 | 0.049573 | 120.6 | 1.56 | 0.1219 | -0.020918 | 0.1753735 |
| (Standardized Latitude-0.00571)*(Standardized Longitude+0.01526) | -0.079106 | 0.0398167 | 119.0 | -1.99 | **0.0492\*** | -0.157948 | -0.000265 |

Fixed Effects Tests
| Source | Nparm | DFNum | DFDen | F Ratio | Prob > F |
| --- | --- | --- | --- | --- | --- |
| Standardized Latitude | 1 | 1 | 118.5 | 4.5384948 | **0.0352\*** |
| Standardized Longitude | 1 | 1 | 114.5 | 3.0873203 | 0.0816 |
| Standardized Altitude | 1 | 1 | 120.6 | 2.4269273 | 0.1219 |
| Standardized Latitude*Standardized Longitude | 1 | 1 | 119.0 | 3.9472082 | **0.0492\*** |

Random Effects Covariance Parameter Estimates
| Variance Component | Var Ratio | Estimate | Std Error | 95% Lower | 95% Upper | Wald p-Value | Pct of Total |
| --- | --- | --- | --- | --- | --- | --- | --- |
| Block | 0.24534 | 4.278413 | 2.3970861 | -0.419789 | 8.9766154 | 0.0743 | 19.334 |
| Genotype | 0.0236 | 0.4116288 | 1.0173377 | -1.582316 | 2.405574 | 0.6858 | 1.860 |
| Residual |  | 17.438679 | 1.6353473 | 14.629984 | 21.146352 |  | 78.806 |
| Total |  | 22.128721 | 2.7429133 | 17.602258 | 28.66712 |  | 100.000 |

Fixed Effects Parameter Estimates
| Term | Estimate | Std Error | DFDen | t Ratio | Prob>\|t\| | 95% Lower | 95% Upper |
| --- | --- | --- | --- | --- | --- | --- | --- |
| Intercept | 3.2161256 | 0.7315537 | 8.0 | 4.40 | **0.0023\*** | 1.530599 | 4.9016521 |
| Standardized Latitude | 0.1542141 | 0.3132987 | 115.1 | 0.49 | 0.6235 | -0.466364 | 0.7747926 |
| Standardized Longitude | 0.1245949 | 0.2406635 | 101.5 | 0.52 | 0.6058 | -0.35279 | 0.6019794 |
| Standardized Altitude | 0.0987799 | 0.3102289 | 108.5 | 0.32 | 0.7508 | -0.516118 | 0.7136781 |

Fixed Effects Tests
| Source | Nparm | DFNum | DFDen | F Ratio | Prob > F |
| --- | --- | --- | --- | --- | --- |
| Standardized Latitude | 1 | 1 | 115.1 | 0.2422875 | 0.6235 |
| Standardized Longitude | 1 | 1 | 101.5 | 0.2680279 | 0.6058 |
| Standardized Altitude | 1 | 1 | 108.5 | 0.1013849 | 0.7508 |

**Random Effects Covariance Parameter Estimates**
| Variance Component | Estimate | Std Error | 95% Lower | 95% Upper | Wald p-Value |
| --- | --- | --- | --- | --- | --- |
| Block | 0.0392746 | 0.02458 | -0.008901 | 0.0874504 | 0.1101 |
| Genotype | 0.0852363 | 0.0260455 | 0.0341881 | 0.1362845 | **0.0011\*** |

**Fixed Effects Parameter Estimates**
| Term | Estimate | Std Error | DFDen | t Ratio | Prob>\|t\| | 95% Lower | 95% Upper |
| --- | --- | --- | --- | --- | --- | --- | --- |
| Intercept | 3.1241504 | 0.0774021 | 9.4 | 40.36 | <.0001* | 2.9501401 | 3.2981607 |
| Standardized Latitude | -0.088531 | 0.0510238 | 115.9 | -1.74 | 0.0854 | -0.189591 | 0.0125286 |
| Standardized Longitude | 0.0519267 | 0.0406299 | 112.3 | 1.28 | 0.2039 | -0.028574 | 0.1324277 |
| Standardized Altitude | -0.111927 | 0.0508265 | 119.7 | -2.20 | **0.0296\*** | -0.212563 | -0.011292 |

**Fixed Effects Tests**
| Source | Nparm | DFNum | DFDen | F Ratio | Prob > F |
| --- | --- | --- | --- | --- | --- |
| Standardized Latitude | 1 | 1 | 115.9 | 3.0105466 | 0.0854 |
| Standardized Longitude | 1 | 1 | 112.3 | 1.6333927 | 0.2039 |
| Standardized Altitude | 1 | 1 | 119.7 | 4.8494503 | **0.0296\*** |

**Supplementary Section 5. Table 11.**
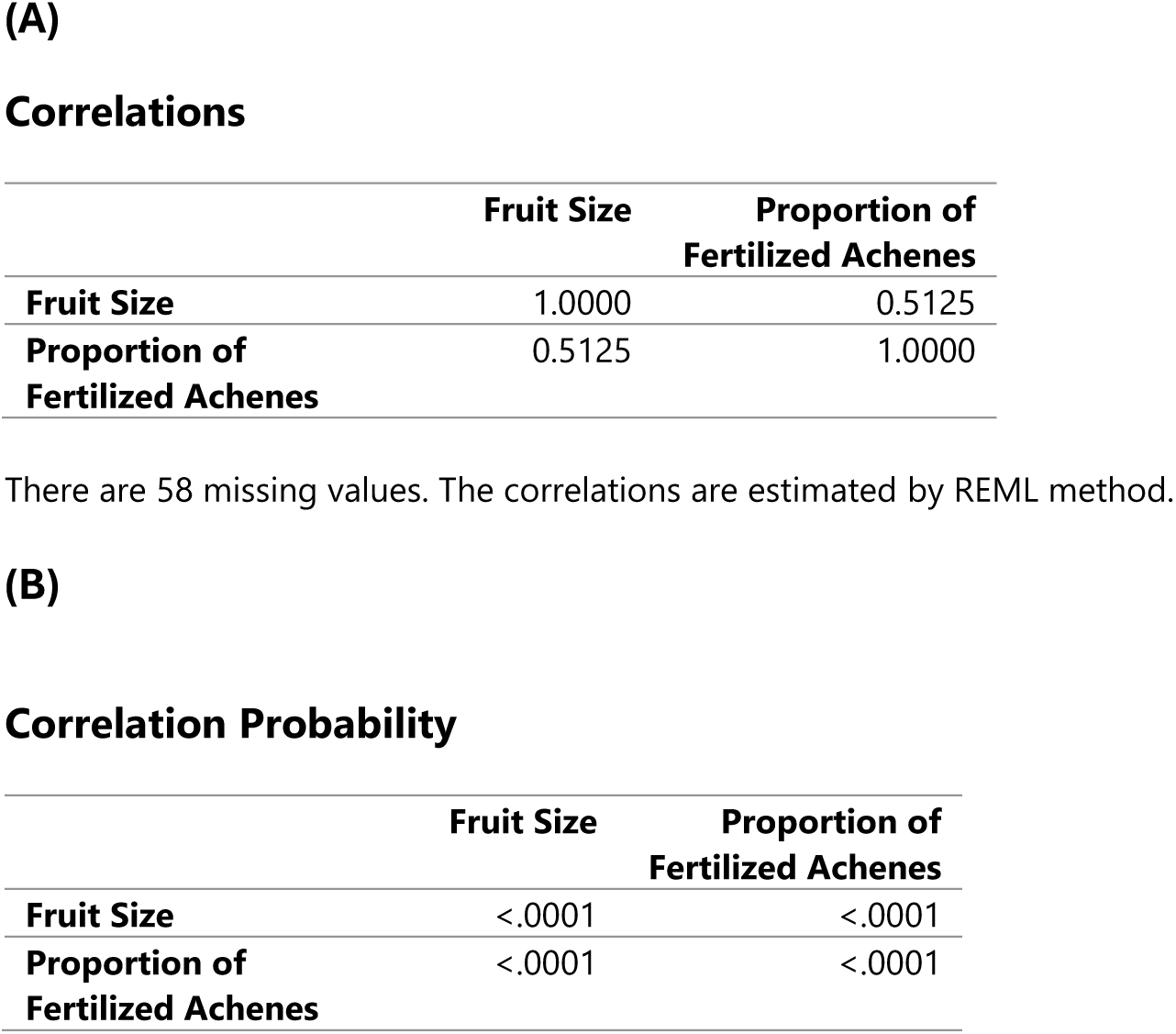
Pearson correlation output between fruit size and the proportion of fertilized achenes. **(A)** correlations coefficients and **(B)** correlation probability

|  | **Fruit Size** | **Proportion of**<br>**Fertilized Achenes** |
| --- | --- | --- |
| **Fruit Size** | 1.0000 | 0.5125 |
| **Proportion of**<br>**Fertilized Achenes** | 0.5125 | 1.0000 |

|  | **Fruit Size** | **Proportion of**<br>**Fertilized Achenes** |
| --- | --- | --- |
| **Fruit Size** | <.0001 | <.0001 |
| **Proportion of**<br>**Fertilized Achenes** | <.0001 | <.0001 |

**Supplementary Section 5. Fig. 1.**
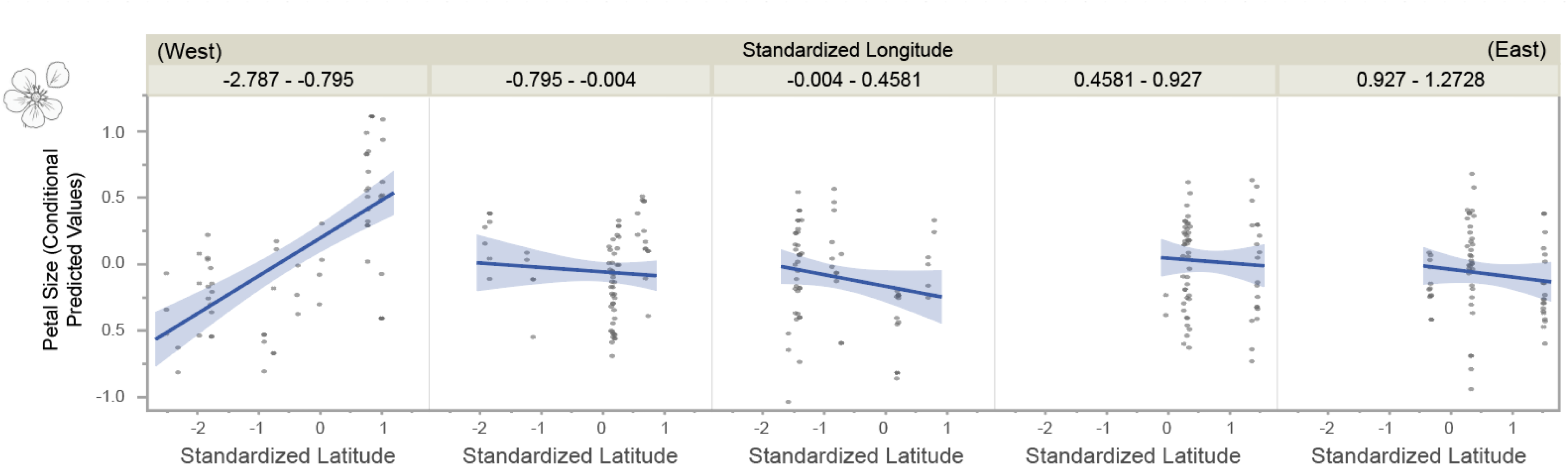
Latitudinal–longitudinal interaction in petal size for *Fragaria vesca*. Petal size (used as a proxy for pollinator attraction) increased strongly from south to north in western Europe (left panel), resulting in the largest petals in the northwest and the smallest in the southwest. In contrast, this relationship was weaker and slightly reversed in eastern Europe, with marginally larger petals toward the southeast. Each point represents a ramet from 121 genotypes. Latitude and longitude were standardized (mean ± SD in the original units: 57.6 ± 8.12° and 10.6 ± 12.14°, respectively). The y-axis shows conditional predicted values of petal size from the model correcting for flower position (see **Supplementary Section 5**, Table 1).

